# IntelliProfiler 2.0: An integrated R pipeline for long-term home-cage behavioral profiling in group-housed mice using eeeHive 2D

**DOI:** 10.64898/2026.02.10.705044

**Authors:** Shohei Ochi, Masashi Azuma, Iori Hara, Hitoshi Inada, Kento Takabayashi, Noriko Osumi

**Affiliations:** Department of Developmental Neuroscience, Tohoku University Graduate School of Medicine, Sendai, Miyagi, 980-8575, Japan; Department of Developmental Neuroscience, Tohoku University School of Medicine, Sendai, 980-8575, Japan; Headquarters for Co-Creation Strategy, National Institutes of Natural Sciences, NINS, Minato-ku, Tokyo, 105-0001, Japan; Kagoshima Seikyo General Hospital, Kagoshima, Kagoshima, 891-0141, Japan

**Author notes:** SO and MA are equally contributed.

**Keywords:** RFID tracking, Group-housed mice, Behavioral phenotyping, Travel distance, Social interaction, R programming

## Abstract

**Background:** Long-term home-cage monitoring is essential to quantify spontaneous locomotor and social behaviors in group-housed mice, but analysis of high-density RFID tracking data remains a barrier to reproducibility.

**New methods:** We developed *IntelliProfiler 2*.*0*, a fully R-based pipeline tailored to the eeeHive 2D floor-mounted RFID array. The workflow performs data import from text logs, preprocessing, coordinate reconstruction, missing-value handling, feature extraction, statistical testing, and visualization in a single environment. Behavioral metrics include travel distance, close contact ratio (CCR), and a newly implemented inter-individual distance metric.

**Results:** In four-day recordings of group-housed C57BL/6J mice (8 males and 8 females), IntelliProfiler 2.0 captured circadian phase-dependent locomotion and proximity patterns and reproduced sex-dependent differences consistent with prior analyses while incorporating updated hardware specifications. Radar-chart summaries enabled intuitive comparison of multidimensional behavioral profiles and inter-individual variability across light/dark phases.

**Comparison with existing methods:** Compared with IntelliProfiler 1.0 and multi-tool workflows, IntelliProfiler 2.0 consolidates analysis into a single, script-based R pipeline, reducing operational complexity and improving reproducibility. The updated implementation supports recent manufacturer-driven changes, including antenna renumbering and multi-USB data export.

**Conclusions:** IntelliProfiler 2.0 provides a reproducible, extensible framework for high-throughput behavioral phenotyping of group-housed mice and is scalable across hardware configurations, including simplified single-board recordings.

**Highlights:**

- End-to-end R pipeline for eeeHive 2D floor-based RFID tracking analysis
- Standardized setup with comprehensive manuals and protocols
- Inter-individual distance metric to quantify group spatial structure
- Circadian- and sex-dependent behavioral profiling in group-housed mice
- Radar-charts summarize multidimensional behavioral profiles and variability

## 1. Introduction

Quantitative behavioral phenotyping of group-housed rodents is central to neuroscience, enabling assessment of spontaneous locomotion, social proximity, circadian organization, and disease-relevant phenotypes under conditions that better approximate everyday living. However, many standard assays (e.g., open field, elevated plus maze, or three-chamber tests) are brief and require handling or transfer to novel arenas, which can induce stress and experimenter-dependent variability (Georgiou et al., 2022; Hurst and West, 2010; Sorge et al., 2014). These constraints motivate long-term, semi-naturalistic home-cage monitoring that captures behavior continuously and with minimal disturbance (Grieco et al., 2021; Jabarin et al., 2022).

Radio-frequency identification (RFID)-based systems provide a practical route to long-term tracking of multiple animals in a shared cage. High-resolution floor-mounted RFID arrays, such as eeeHive 2D (Phenovance LLC, Japan), record spatiotemporal positions of individually tagged animals and enable quantification of locomotor activity and proximity-based social behavior in freely interacting groups (Lipp et al., 2024). To analyze such high-density RFID data, we previously developed IntelliProfiler (version 1.0), a workflow tailored to the original RFID floor plate system, which quantified travel distance and a proximity-based index, the close contact ratio (CCR).

Despite its utility, IntelliProfiler 1.0 relied on multiple separate software tools, (e.g., R, Python, spreadsheets, and commercial plotting software, which increased friction for users and complicated reproducible execution across laboratories. In addition, manufacturer-driven updates to eeeHive hardware (including antenna numbering and multi-USB data export) required systematic updates to the analysis workflow.

Here we present IntelliProfiler 2.0, a fully integrated, R-only pipeline that performs data import, preprocessing, positional reconstruction, feature extraction, statistical testing, and visualization within a single environment. We demonstrate analytical consistency using new recordings from group-housed male and female C57BL/6 mice and introduce inter-individual distance as an additional metric to quantify spatial relationships among animals. We further provide radar-chart visualization to summarize multidimensional behavioral profiles and day-by-day dynamics. Together, IntelliProfiler 2.0 provides a standardized and scalable framework for long-term behavioral analysis of group-housed mice using high resolution floor RFID tracking.

## 2. Materials and Methods

### 2.1 Animals and ethics approval

C57BL/6J mice (7-week-old) were obtained from CLEA Japan and maintained in the animal facility at Tohoku University Graduate School of Medicine. All procedures below were approved by the Ethics Committee for Animal Experiments at Tohoku University Graduate School of Medicine (approval ID: 2021MdA-020-15).

### 2.2 RFID tag implantation

Mice were anesthetized using isoflurane (5.0% for induction, 2.0% for maintenance) delivered via an induction chamber and facemask (**Fig. 1A**). RFID tags (7 mm × 1.25 mm; Phenovance LLC, Japan) were subcutaneously implanted into the abdominal region using sterile preloaded injectors (**Fig. 1B**). After implantation, tag IDs were verified using an HF (high frequency) RFID reader (Phenovance LLC, Japan) (**Fig. 1C**). Mice were placed on a heating plate (Tokyo Garasu Kikai Co., Ltd., Japan) maintained at 37 °C for approximately 5 minutes and monitored until fully awake (**Fig. 1D**). A detailed surgical protocol is provided in **Supplementary Information Materials** and **Supplementary Movie 1**. Recordings were conducted when mice were 8-9 weeks old.

**Figure 1.**
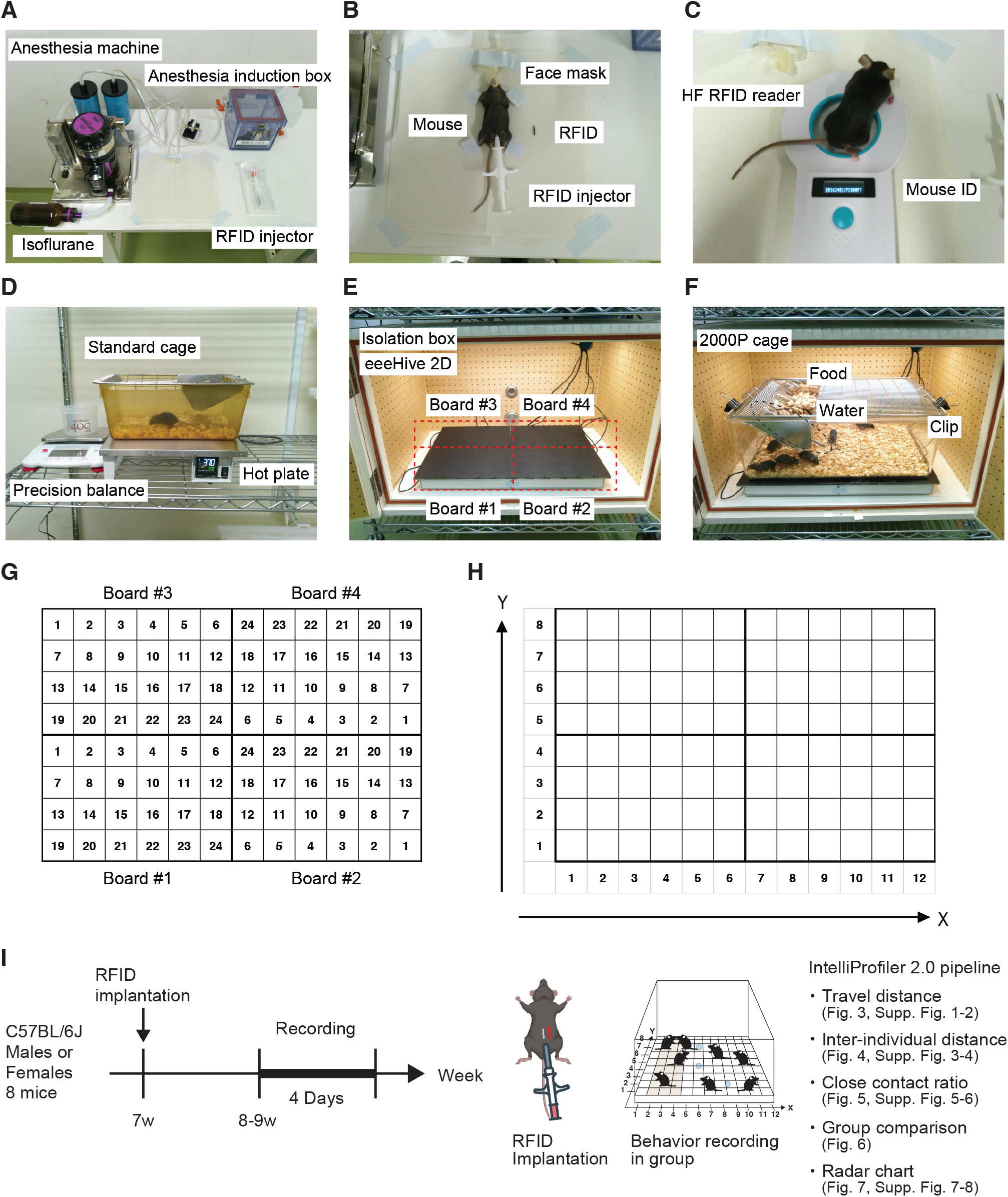
Overview of the eeeHive 2D high-resolution antenna array and the IntelliProfiler 2.0 workflow for group behavior monitoring. (A-F) Photographs of the experimental setup. (A) Anesthesia system and implantation environment. (B) RFID tag implantation under anesthesia. (C) Tag ID verification using a high-frequency (HF) RFID reader. (D) Postoperative warming and recovery. (E) eeeHive 2D system with soundproof enclosure. (F) Group behavior recording in a 2000P cage. (G) Layout and antenna numbering of the four eeeHive 2D boards. (H) Conversion of antenna positions to XY coordinates (X1–12, and Y1–8) in IntelliProfiler 2.0. (I) Experimental timeline. RFID tags were implanted in 7-week-old C57BL/6 mice. Group behavior was recorded for 4 consecutive days at 8–9 weeks old. Abbreviations: RFID, radio-frequency identification.

### 2.3 Hardware and recording environment

The eeeHive 2D RFID antenna board (Phenovance LLC, Japan) consists of 24 antenna tiles (5 cm x 5 cm) arranged in a 6 x 4 layout. For the primary experiments, four boards were placed adjacent to cover the floor of a standard mouse home cage, yielding 96 antenna tiles and a total coverage 60 cm x 40 cm (**Fig. 1E**). A 2000P mouse home cage (58 cm × 38 cm × 21.5 cm; Tecniplast, Italy) with a lid containing food and water bottles (CLEA Japan) was positioned directly above the RFID floor array inside a sound-attenuating chamber (Phenovance LLC, Japan) (**Fig. 1F**). Animals were maintained on a 12:12 hour light/dark cycle (lights on at 08:00, off at 20:00) was maintained using a programmable outlet timer (Levex, Japan). Mice were introduced to the recording environment at 8:00 on Day 1 (light phase) and recorded continuously for four consecutive days. A complete list of equipment and housing components is provided in **Supplementary Table 1**.

### 2.4 Manufacturer updates to eeeHive specifications and implications for analysis

Since the initial IntelliProfiler 1.0 report (Ochi et al., 2026), the commercially available RFID floor plate system has undergone manufacturer-driven updates that affect how raw data are labeled and exported. IntelliProfiler 2.0 was updated to accommodate these changes.

1. **Antenna numbering**. The original multi-board configuration used a vertically increasing antenna numbering scheme across the four-board array (previous 1–96 mapping; Ochi et al., 2026). The updated eeeHive 2D specification applies a left-to-right numbering scheme within each board (1-24 per board, **Fig. 1G**).
2. **USB architecture and data-export**. In the earlier configuration, data were exported through a single USB connection from a primary board, with additional boards linked via ribbon cables (Ochi et al., 2026). In the updated specification, each RFID board connects directly to the host computer via its own USB port, and data from each board are exported independently through multiple USB connections.

### 2.5 Software implication

IntelliProfiler 2.0 is implemented entirely in R (R Core Team, 2024) and executed in RStudio. R (version 4.5.0) was used for preprocessing, statistical analysis, and visualization. Required packages and script dependencies are listed in **Supplementary Tables 2 and 3**. Custom scripts are available in a public GitHub repository (see “**Code Availability**”).

### 2.6 IntelliProfiler 2.0 analysis pipeline

Raw log files exported via terminal loggings (Tera Term) contain timestamps, board IDs, antenna IDs (1-24), and transponder IDs. IntelliProfiler 2.0 imports and merges the four-board outputs and converts the manufacturer-specific antenna indices into a unified two-dimensional coordinate matrix (X1–12, Y1–8; **Fig. 1H**). Data are then binned to 1-s resolution to generate time-aligned trajectories for each animal. Missing reads are identified and handled during preprocessing (with optimal imputation as described in the **Supplementary Information**). An overview of the experimental workflow is shown in **Fig. 1I**. The pipeline comprises five steps (**Figure 2**): (1) data collection, (2) preprocessing and positional reconstruction, (3) feature extraction, (4) group-level comparison, and (5) visualization (including using radar charts). IntelliProfiler 2.0 pipeline represents an updated version of our previous IntelliProfiler 1.0 workflow (Ochi et al., 2026). All procedures were modified to accommodate the redesigned eeeHive2D hardware (Lipp et al., 2024, Phenovance LLC, Japan), including the revised antenna layout and the multi-channel USB configuration.

**Figure 2.**
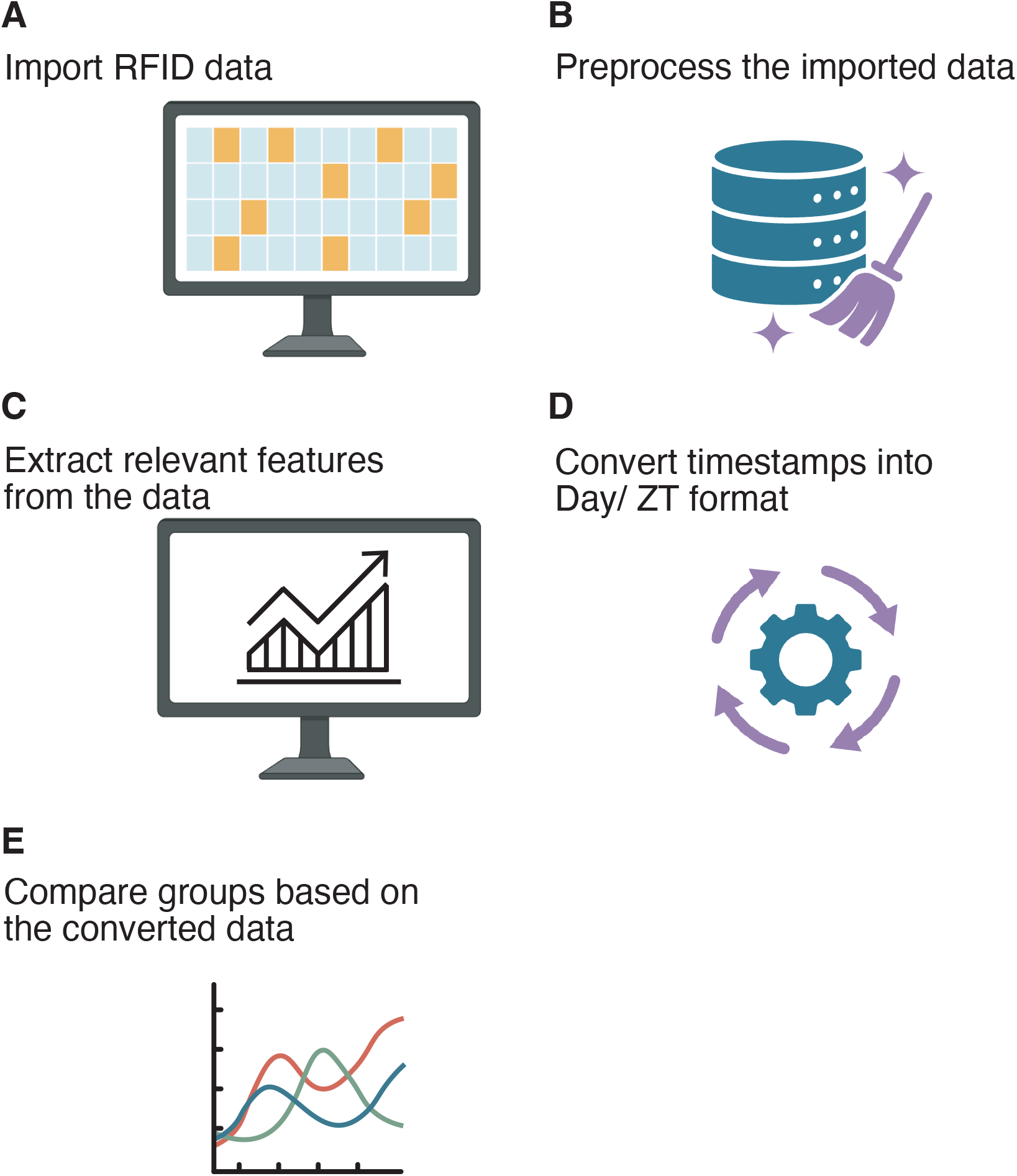
Overview of the IntelliProfiler 2.0 pipeline. IntelliProfiler 2.0 processes raw RFID data collected from the eeeHive 2D floor array in five steps. (A) Import text-based log files. (B) Preprocess and merge data and reconstruct positional time-series matrices. (C) Compute core metrics (travel distance, inter-individual distance, and CCR). (D) Convert timestamps to biologically meaningful units (Day and ZT) and generate standardized tables for phase-based analyses. (E) Statistically compare groups (e.g., sexes, or conditions) and visualize results. Schematic elements were created with BioRender. Abbreviations: CCR, close contact ratio, ZT, Zeitgeber Time.

Behavioral metrics were identified as follows. Travel distance was computed as the cumulative distance traveled across time bins based on the Euclidean distance between consecutive XY positions. Inter-individual distance was calculated as the Euclidean distance between XY positions for each mouse pair at each time bin and summarized as a time series for each pair. Close contact ratio (CCR) was operationally defined as two mice occupying the same grid cell or immediately adjacent cell, and CCR was calculated as the percentage of time in close contact within each time window (Fig. 5A).

To facilitate circadian analyses, timestamps were converted to Zeitgeber time (ZT) with ZT0 defined as lights on (08:00). Metrics were summarized in 1-h windows (ST), in 12-h light/dark windows, and as dark/light ratio on each day.

### 2.7 Statistics

Group differences in travel distance, inter-individual distance, and CCR were evaluated using two-way analysis of variance (ANOVA) followed by Tukey’s *post hoc* tests. Analyses were performed (i) for each ZT hour and (ii) for each 12-hour light/dark phase. Statistical significance is defined as follows: ^*^*p* < 0.05, ^**^*p* < 0.01, ^***^*p* < 0.001. All analyses and visualizations were performed in R (version 4.5.0). Full statistical procedures are provided in **Supplementary Information**.

## 3. Results

### 3.1. Group behavioral recording and IntelliProfiler 2.0 workflow

Photographs of the experimental setups and recording environment are shown in **Fig. 1A–F**. Male and female C57BL/6J mice were implanted with RFID tags (**Fig. 1B, I**; **Supplementary Movie 1**) and allowed to recover for at least a one week. Group behavior was recorded continuously for four days in the home-cage environment (**Fig. 1F, I**). Raw data (time stamps, board ID, antenna ID, and transponder ID) were exported from four eeeHive 2D boards that tiled the entire cage floor (**Fig. 1F–G**). These time-series data were merged, converted to XY coordinates (X1–12, Y1–8), and downsampled to 1-s resolution using IntelliProfiler 2.0 pipeline (**Fig. 1H**). Body weight was measured before and after monitoring to enable exploratory correlation analyses with behavioral parameters (**Fig. 1I**). The updated workflow was developed to accommodate the lates hardware version (eeeHive 2D v1.2).

### 3.2. Temporal and sex-dependent patterns of locomotor and proximitry-based behaviors

Across four days of continuous monitoring, IntelliProfiler 2.0 identified reproducible temporal structure in locomotor activity and a proximity-based measures in both sexes (**Supplementary Fig. 1–6**, and representative examples are shown in **Fig. 3–6**). Travel distance and inter-individual distance were transiently elevated immediately after cage entry and decreased over the first hours of recording (**Fig. 3A, B, G, H; Fig. 4A, B, G, H; Supplementary Fig. 1A, 2A, 3A, 4A**). When summarized in 12-h windows, both measures were higher during the dark phase than the light phase (**Fig. 3C, D, I, J; Fig. 4C, D, I, J; Supplementary Fig. 1B, 2B, 3B, 4B**). Dark/light ratios suggested relatively stable circadian organization across days, with greater day-to-day variability in females for some measures (**Fig. 2E, F, K, L; Fig. 3E, F, K, L; Supplementary Fig. 1C, 2C, 3C, 4C**).

**Figure 3.**
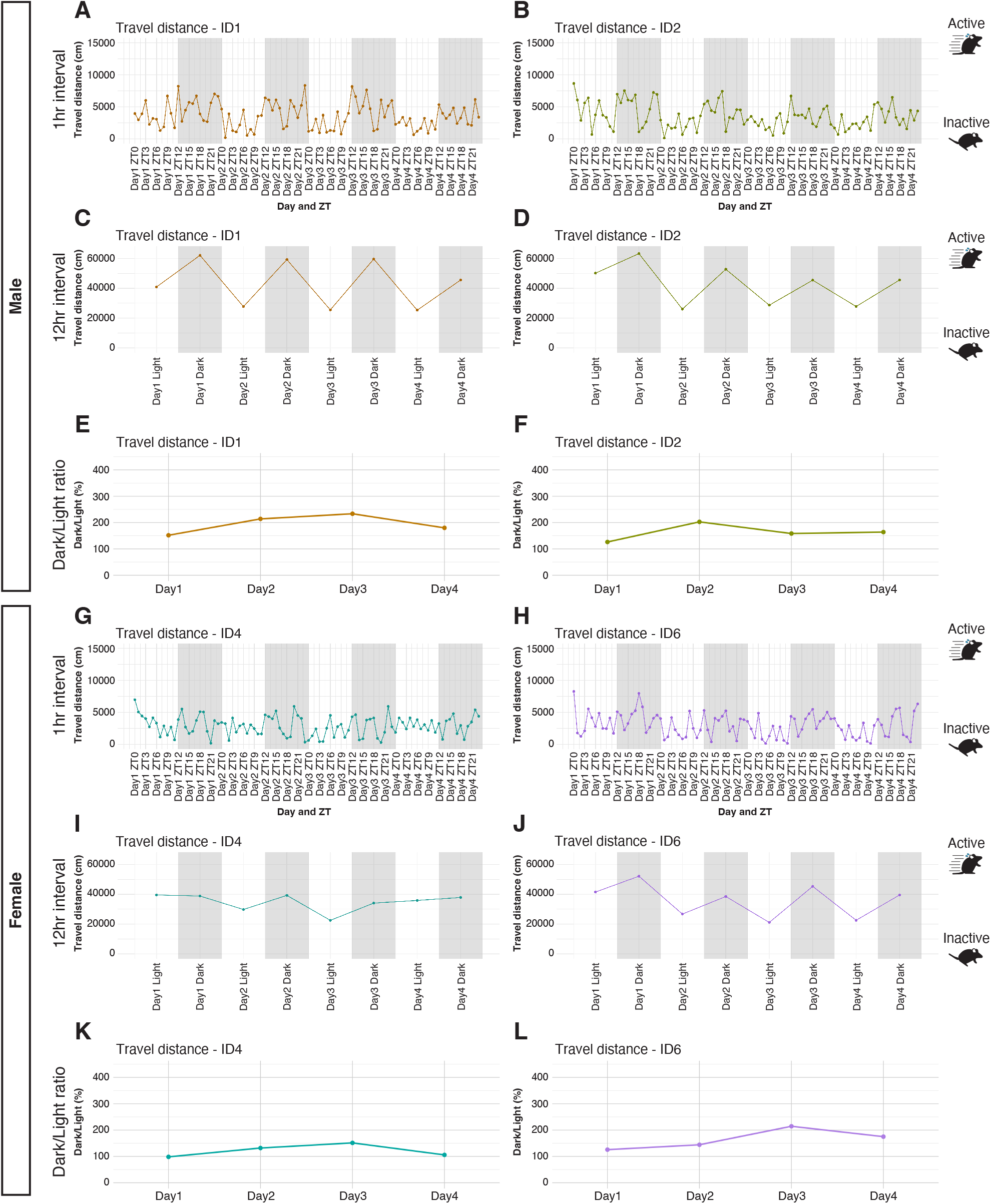
Temporal patterns of travel distance in individual mice using the IntelliProfiler 2.0. (A-L) Line plots show cumulative travel distance over time for individual male and female mice (n=8 per sex). Two representative males (A-F) and two representative females (G-L) are shown. Travel distances are summarized in 1-hour windows (A, B, G, H), 12-hour light/dark windows (C, D, I, J), and as dark/light ratios per each day (E, F, K, L). Abbreviation: ZT, zeitgeber time.

**Figure 4.**
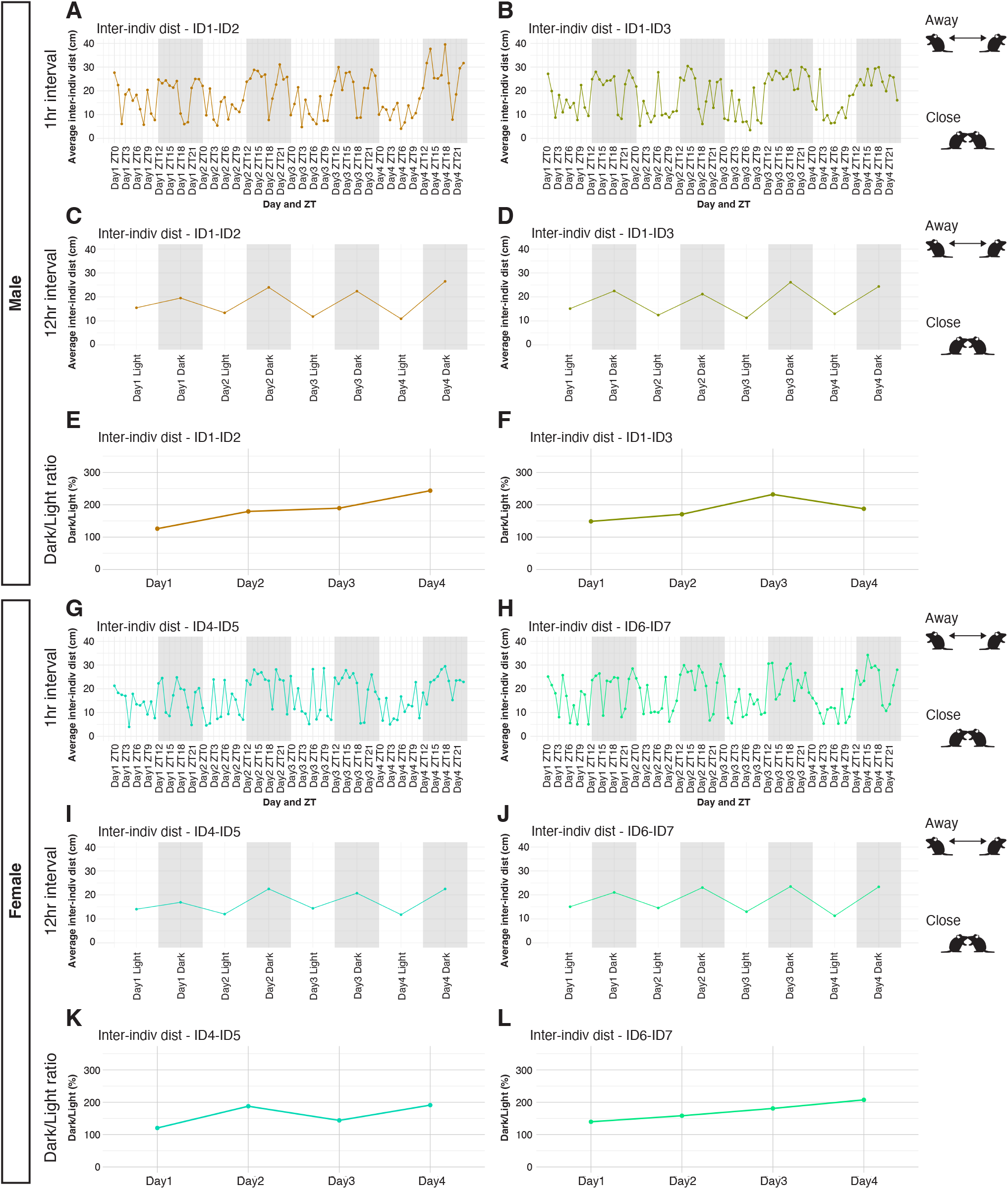
Temporal patterns of inter-individual distance between mouse pairs using IntelliProfiler 2.0. (A-L) Line plots show average inter-individual distance over time for male and female mouse pairs (n=8 per sex; 28 possible pair combinations per group). Two representative male pairs (A-F) and two representative female pairs (G-L) are shown. Inter-individual distances are summarized in 1-hour windows (A, B, G, H), 12-hour light/dark windows (C, D, I, J), and as dark/light ratios per each day (E, F, K, L). Abbreviation: ZT, zeitgeber time.

In contrast, CCR showed an opposite phase dependency, with higher values during the light phase and larger fluctuations across days in some female pairs (**Fig. 5B–M; Supplementary Fig. 5-6**). Group comparisons revealed higher cumulative travel distance in males at multiple time points, particularly during the light phase on Days 3– 4 (**Fig. 6A, D, G, J**), while inter-individual distance showed no consistent sex-dependent bias (**Fig. 6B, E, H, K**). CCR tended to be higher in females, especially during the light phase on Days 3–4 (**Fig. 6C, F, I, L**). Heatmaps of individual- and pair-level values highlighted substantial within-sex variability while supporting these overall trends (**Fig. 6D–F, J–L**).

**Figure 5.**
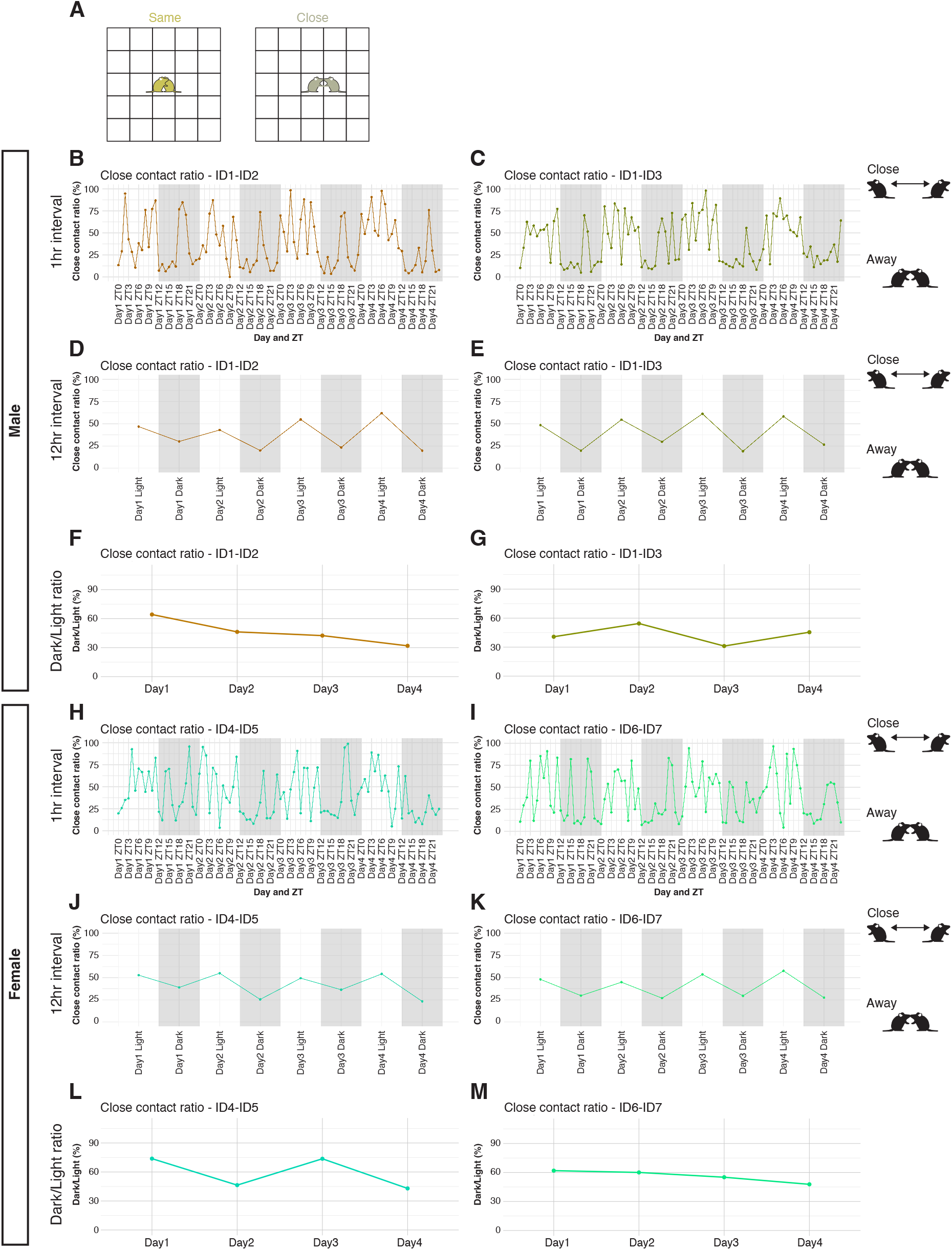
Temporal patterns of close contact ratio (CCR) between mouse pairs using IntelliProfiler 2.0. (A)Close contact was defined as two mice occupying the same grid cell (yellow) or in immediately adjacent cells (olive). (B-M) Line plots show CCR (percentage of time in close contact) over time for male and female mouse pairs (n=8 per sex; 28 possible pair combinations per group). Two representative male pairs (B-G) and two representative female pairs (H-M) are shown. CCRs are summarized in 1-hour windows (B, C, H, I), 12-hour light/dark windows (D, E, J, K), and as dark/light ratio per day (F, G, L, M). Abbreviation: ZT, zeitgeber time.

**Figure 6.**
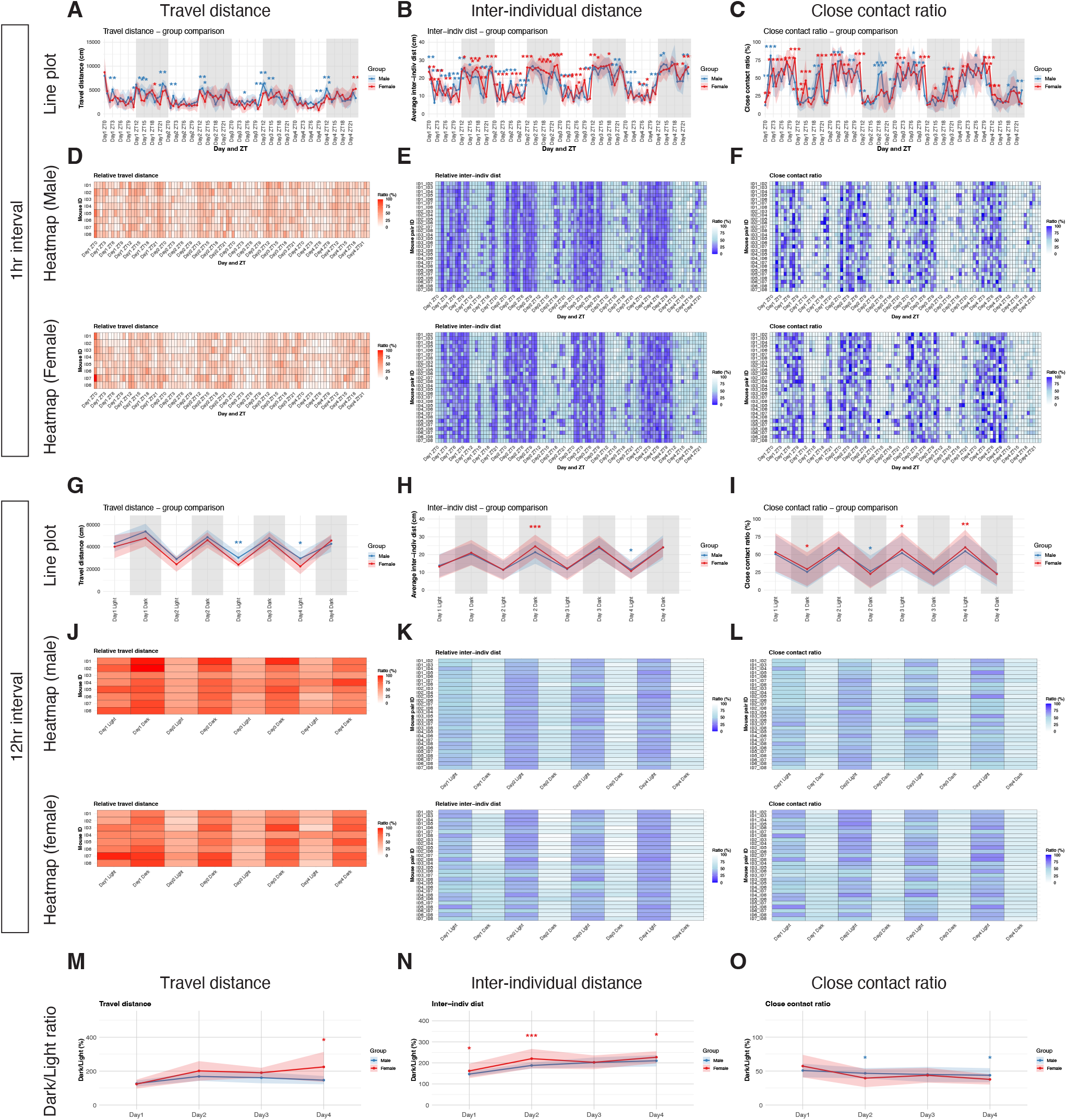
Group comparison of locomotor activity and proximity-based behavior in male and female mice using IntelliProfiler 2.0. (A–O) Comparisons between male and female groups (n=8 per sex) across four consecutive days. Panels show cumulative travel distance (A, D, G, J, M), average inter-individual distance (B, E, H, K, N), and CCR (C, F, I, L, O). Metrics are summarized in 1-hour windows (A-F), 12-hour light/dark windows (G-L), and as dark/light ratios per day (M-O). Line plots are shown in (A-C, G-I, M-O), and heatmaps in (D-F, J-L).Statistical testing followed the approach described in Methods; significance is indicated as \**p* < 0.05, \*\**p* < 0.01, \*\*\**p* < 0.001. Male and female data are shown in blue and red, respectively; shaded ribbons indicate standard deviation (SD). Abbreviations; CCR, close contact ratio.

Circadian indices further confirmed sex-dependent differences: females showed higher dark/light ratios for travel distance on Day 4 and for inter-individual distance on Days 1, 2, and 4 (**Fig. 6M, N**), whereas males exhibited higher dark/light CCR ratios on Days 2 and 4 (**Fig. 6O**). These analyses collectively demonstrate that the IntelliProfiler 2.0 pipeline captures nuanced temporal structures of locomotor and social behaviors, enabling quantitative comparisons of sex-dependent and circadian behavioral profiles in group-housed mice.

### 3.3 Radar chart visualization of behavioral parameters

To provide an intuitive overview of multidimensional behavior, we implemented radar chart visualization using six normalized parameters: travel distance (light), travel distance (dark), inter-individual distance (light), inter-individual distance (dark), CCR (light), and CCR (dark) referring our previous behavioral evaluation (Yoshizaki et al., 2017). Values from all 16 mice were first scaled to a 0–1 range for each parameter, and the male group mean averaged across Days 1–4 was the set to 1.0 as a reference, allowing relative comparison across parameters and animals.

Individual-level radar charts (averaged across Days 1–4) revealed heterogeneity in behavioral profiles, including a subset of females with relatively higher light-phase CCR and/or higher dark-phase inter-individual distance (**Fig. 7A–B**). Day-by-day radar charts indicated that profiles were largest and most variable on Day 1, immediately after introduction into the recording environment and became progressively more stable by Days 3-4 (**Supplementary Fig. 7A–B**).

**Figure 7.**
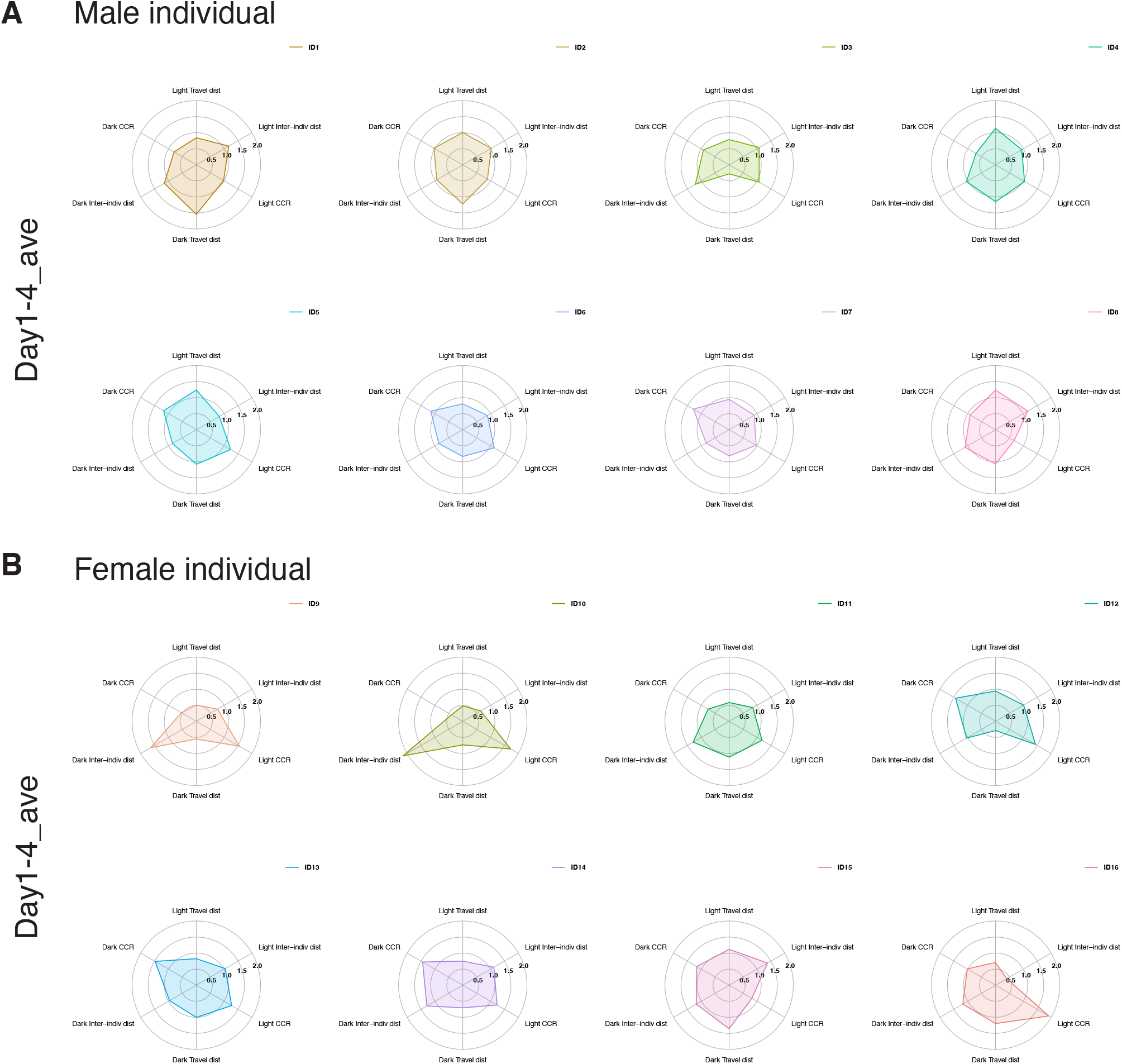
Individual-level radar chart profiles of locomotor activity and proximity-based behavior using IntelliProfiler 2.0. (A, B) Radar charts summarizing individual behavioral profiles for male (A) and female groups (n=8 per sex) from Day 1 to Day 4. Each chart represents the average profile of single mouse over the indicated period. Parameters include travel distance, inter-individual distance, and CCR, each quantified for light and dark phases. Abbreviations; CCR, close contact ratio.

We next evaluated group-level sex differences and day-to-day dynamics using radar charts constructed from group means (**Fig. 8**). The radar plots summarize behavioral profiles for each recording day (Day 1-4) and for the overall mean across Days 1–4. Across the recording period, clear sex- and time-dependent patterns emerged. On Day 3, females exhibited higher light-phase CCR, while males showed higher light-phase travel distance. On Day 4, males displayed higher light-phase travel distance and higher light-phase inter-individual distance, whereas females again showed higher light-phase CCR. Consistent trends were also evident in the 4-day average: males exhibited higher travel distance during the light phase, whereas females showed higher CCR during the light phase and higher inter-individual distance during the dark phase.

**Figure 8.**
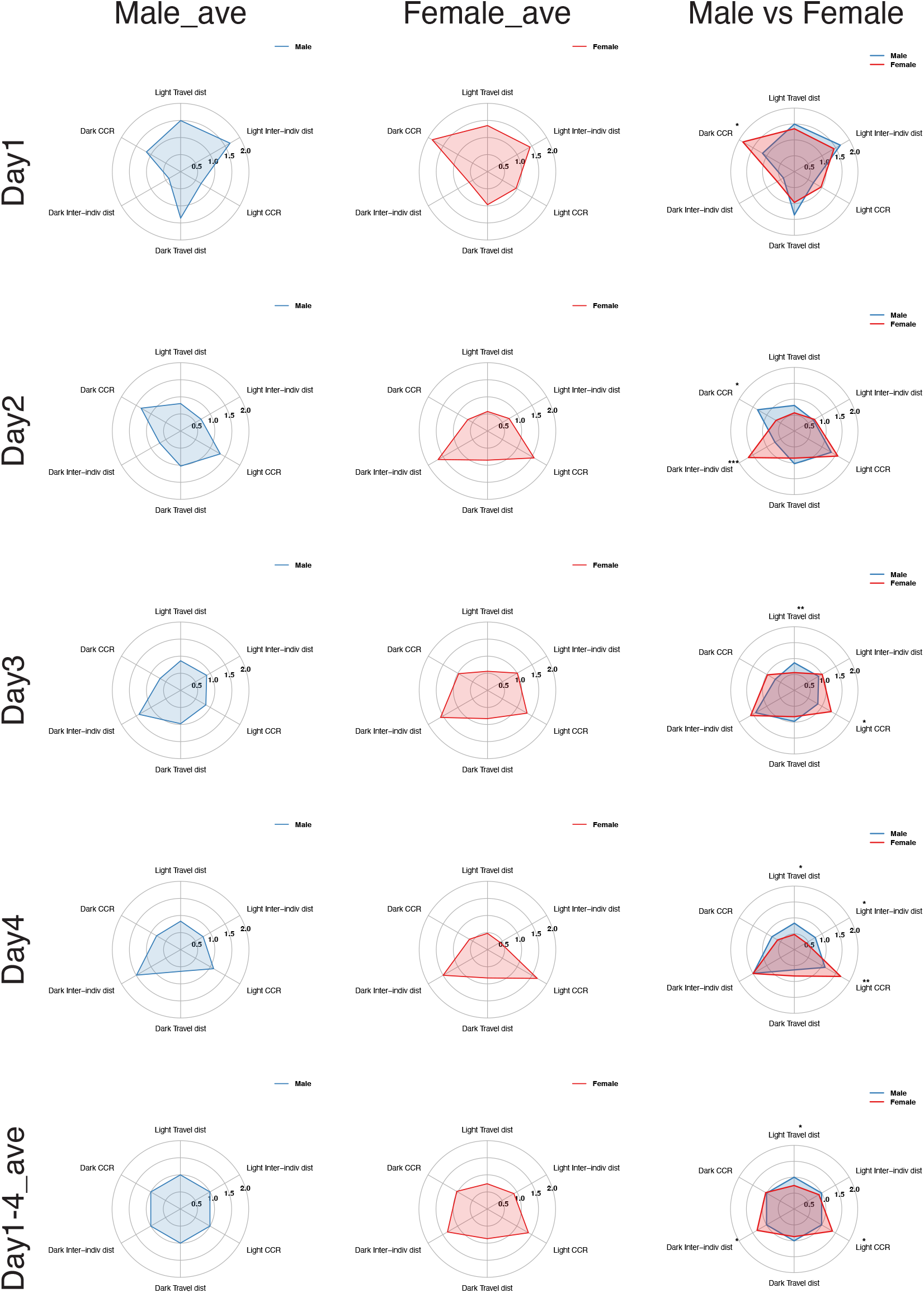
Group-level radar chart comparison of behavior profiles between sexes using IntelliProfiler 2.0. Mean radar charts for male group (left) and female group (middle) groups (n=8 per sex) from Day 1 to Day 4 and for the average across Days 1–4. Parameters include travel distance, inter-individual distance, and CCR, each quantified separately for the light and dark phases. For each behavioral parameter, male and female groups were compared using Student’s *t*-test at each corresponding time window. Statistical significance is indicated as ^*^*p* < 0.05, ^**^*p* < 0.01, ^***^*p* < 0.001. Abbreviations; CCR, close contact ratio.

Collectively, these results suggest that behavioral profiles are comparatively labile immediately after cage introduction but become progressively more organized over Days 1-4. As monitoring continued, males tended to increase locomotor activity, whereas females exhibited increasingly prominent proximity-based interaction patterns. Thus, radar charts provide an intuitive visualizing of multidimensional behavioral signatures, facilitating comparisons of temporal trajectories and sex differences across multiple parameters.

## 4. Discussion

### 4.1 Advantages of an end-to-end R workflow

IntelliProfiler 2.0 consolidates the full analysis workflow for high-resolution RFID tracking into a single R/RStudio environment. Compared with the multi-tool IntelliProfiler 1.0 workflow (Ochi et al., 2026), this integration reduced friction, improves traceability of analytical decisions, and facilitates reproducible execution across users and laboratories. Because the complete pipeline is script-based and openly shared, users can audit, modify, and extend the analysis to match experimental needs while maintaining a documented computational record.

### 4.2 Capturing temporally structured locomotor and proximity-based phenotypes

Using four-day recording from group-housed mice, IntelliProfiler 2.0 quantified exploration/habituation dynamics after cage introduction and robust circadian organization across locomotor and proximity-based measures (**Fig. 3–7**). Higher travel distance and greater inter-individual spacing during the dark phase, together with higher CCR during the light phase, are consistent with nocturnal activity patterns (Grieco et al., 2021; Jhuang et al., 2010) as well as with previous observation using video analysis and floor-RFID tracking (Endo et al., 2018; Ochi et al., 2026) . IntelliProfiler 2.0 also reproduced sex-dependent differences in locomotion and CCR reported previously (Ochi et al., 2026), supporting analytical consistency across hardware versions.

Inter-individual distance complements CCR by capturing spatial dispersion and group structure beyond binary proximity events. Together, these measures provide a compact summary of group organization over time while remaining interpretable and computationally lightweight.

### 4.3 Visualization for high-dimensional summaries

Radar charts provide a compact visualization to summarize multiple behavioral parameters simultaneously and to compare individuals, days, and groups at a glance. While radar charts do not replace statistical testing, they can be useful for exploratory assessment of behavioral “profiles”, identification of outliers, and communication of multidimensional results to broad audiences.

### 4.4 Scalability, configurations, and practical constraints

The analysis workflow is compatible with different hardware configurations. In addition to the four boards (40 cm × 60 cm) setup used here, we tested a smaller single-board (20 cm × 30 cm; “IP-Single”) to analyze four mice and observed sex-related behavioral differences consistent with the multi-board setup (**Supplementary Fig. 8**). This flexibility may facilitate adaptation in laboratories with limited space of throughput requirements.

### 4.5 Limitations and future directions

Several limitations should be acknowledged. First, RFID-based floor tracking provides horizontal position at tile-level resolution and does not capture posture, fine-grained social actions, or vertical behaviors (e.g., climbing onto the cage lid). Bedding height and cage accessories can also influence detection performance. Second, the current workflow focuses on travel distance, inter-individual distance, and CCR; richer ethological classification will require integration with video tracking or behavior-segmentation approaches. A multimodal framework combining RFID with video-derived pose or interaction features is a promising direction for future IntelliProfiler development. Finally, while IntelliProfiler 2.0 is designed to be general within the eeeHive ecosystem, parameter choices (e.g., windowing, definitions of proximity) should be reported explicitly to support comparison across studies.

The flexibility of the RFID-based framework enables diverse applications, including disease model studies (e.g., autism spectrum disorder and neurodegenerative disease models) and pharmacological interventions (Kubota et al., 2025; Maisterrena et al., 2024; Morito et al., 2025; Watamura et al., 2025). Future work will benefit from leveraging eeeHive 2D together with IntelliProfiler 2.0 as a standardized, scalable approach for long-term monitoring and quantitative comparison of naturalistic group behaviors.

## 5. Conclusion

IntelliProfiler 2.0 is an integrated, R-based pipeline for long-term behavioral analysis of group-housed mice tracked with a high-resolution RFID system. By providing end-to-end processing, standardized metrics (including inter-individual distance and CCR), and intuitive multidimensional visualization, IntelliProfiler 2.0 supports reproducible and scalable home-cage behavioral phenotyping across neuroscience and related fields.

## Acknowledgements

We thank Dr. Toshihiro Endo, Dr. Seico Benner, Prof. Hans-Peter Lipp, and their colleagues for developing the eeeHive 2D RFID floor array. We also thank Dr. Endo for technical advice.

## CRediT authorship contribution statement

**Shohei Ochi**: Conceptualization, Methodology, Animal experiments, Software, Validation, Formal analysis, Investigation, Data curation, Writing – Original draft, Writing – Review & Editing, Visualization, Supervision, Project administration, Funding acquisition. **Masashi Azuma**: Methodology, Software, Formal analysis, Data curation, Visualization, Writing – Review & Editing, Visualization. **Iori Hara**: Methodology, Software, Formal analysis, Data curation, Visualization. **Hitoshi Inada**: Conceptualization, Methodology, Software, Writing – Review & Editing. **Kento Takabayashi**: Methodology, Software. **Noriko Osumi**: Conceptualization, Writing – Review & Editing, Supervision, Project administration, Funding acquisition.

## Code Availability

All scripts used for data analysis are available on GitHub.

- Repository URL: https://github.com/IntelliProfiler/IntelliProfiler2.0

