## Supplementary Information for "IntelliProfiler 2.0: An integrated R pipeline for long-term home-cage behavioral profiling in group-housed mice using eeeHive 2D"

### Materials and Methods

#### S1.1 Detailed protocol for RFID tag implantation

The current methodology is primarily based on our previous article (Ochi et al., 2026), with minor updates. Mice were anesthetized using 5.0% isoflurane (MSD Animal Health, Japan) for induction and 2.0% maintenance, delivered via an isoflurane induction chamber (**Fig. 1A**; SHINFACTORY, Japan) and a facemask (**Fig. 1B**; Bio Research Center, Japan). Once fully anesthetized, mice were placed in a supine position, and their abdominal region was disinfected with a 70% ethanol-soaked Kimwipe (NIPPON PAPER CRECIA). The abdominal skin was gently lifted by pinching to facilitate subcutaneous access. A 7 mm × 1.25 mm RFID tag (Phenovance LLC, Japan), preloaded in a sterile injector, was inserted subcutaneously at a shallow angle from caudal to cranial to minimize the risk of puncturing the abdominal cavity. After the needle was fully advanced, the tag was ejected and stabilized with light pressure during needle withdrawal. The RFID identifier was read using a high-frequency (HF) handheld RFID reader (**Fig. 1C**; Phenovance LLC, Japan) and manually recorded at the time of implantation. Mice were then placed in standard plastic cages (CLEA Japan) on a 37 ℃ warming plate (Tokyo Glass Instruments, Japan) and monitored until fully awake (**Fig. 1D**). After recovery, mice were returned to standard housing conditions, and behavioral recording began after a recovery period of at least one week. Representative photographs are shown in **Figure 1A–D**, and a detailed video protocol is provided in **Supplementary Movie 1**. A comprehensive list of equipment, materials, and animals is provided in **Supplementary Table 1**.

#### S1.2. Data preprocessing

Behavioral data were acquired from the eeeHive2D high-resolution RFID antenna array (Phenovance LCC, Japan; Lipp et al., 2024), which continuously recorded timestamps, board IDs, antenna IDs, and transponder IDs at sub-second resolution. Raw data were exported via Universal Serial Bus (USB) using Tera Term software as plain text log files. Subsequent preprocessing was performed using a custom R script (IP_RawLog_to_PositionID.R; **Supplementary Table 2**), which carried out the following steps:

(1) Coordinate remapping: Antenna board positions (1–96) were converted into a unified X-Y coordinate system spanning X = 1–12 and Y = 1–8, corresponding to the spatial layout of the combined antenna array covering the entire cage floor area;

(2) Time normalization: Data were restructured into a per-second time series matrix containing time, antenna ID, transponder ID, and X-Y coordinates;

(3) Missing data imputation: Occasional gaps in detection were interpolated using a last observation carried forward (LOCF) method;

(4) Data consolidation: Processed data were saved both as a combined Excel file (processed_data_combined.xlsx) and as individual Excel files per mouse ID (e.g., E01624011FC6507F.xlsx), to facilitate downstream behavioral analyses.

The core R pipeline used the following packages: dplyr, tidyr, lubridate, doParallel, openxlsx, readr, purrr, and foreach. Parallel processing was implemented via the doParallel package to accelerate per-ID file generation (**Supplementary Table 3**).

**S1.3. Behavioral feature extraction**

Behavioral features were extracted from the preprocessed position data using a dedicated R script (IP_FeatureCalc_Visualization.R; **Supplementary Table 2**). The script computed key locomotor and social interaction metrics updated from our previously published pipeline (Ochi et al., 2026) with updates for IntelliProfiler 2.0

(1) Travel distance: for each mouse, per-second travel distance was calculated as the Euclidean displacement between consecutive X-Y coordinates:

$$D_{t+1} = \sqrt{(X_{t+1} - X_t)^2 + (Y_{t+1} - Y_t)^2} \times 5\ (cm)$$

The per-second distances were accumulated over time to yield total travel distance across 1-hour intervals.

(2) Inter-individual distances: pairwise Euclidean distances between all possible mouse pairs were calculated at each timepoint as:

$$L_t^{(i,j)} = \sqrt{\left(X_t^{(i)} - X_t^{(j)}\right)^2 + \left(Y_t^{(i)} - Y_t^{(j)}\right)^2} \times 5\ (cm)$$

where *i, j* indicate the identities of individual mice, and *t* is the time point.

(3) Close contact ratio (CCR): CCR was defined as the percentage of time two mice spent in close proximity (<10 cm) within each 1-hour interval:

$$CCR_{ij} = \left(\frac{1}{T}\sum_{t=1}^{T} I\left(D_{ij}(t) < 10\right)\right) \times 100\ (\%)$$

where *i* and j indicate the identities of individual mice, and *t* is the time point (1-second resolution; T=3600 per hour).

The following output files were generated for each experimental session:

- Hourly_movement.xlsx: Total travel distance per mouse, binned in 1-hour intervals (cm/hour);
- Hourly_inter-individual_distance.xlsx: Mean pairwise distances between two mice, calculated per 1-hour interval (cm/hour);
- Close_contact_ratio.xlsx: CCR, representing the percentage of time in proximity per hour (CCR [%/hour]);
- ID_mapping.txt: Mapping of original transponder IDs to shortened IDs used in visualization (e.g., ID1 = E01624011FC6507F);
- movement_hour.pdf: Line plot of hourly travel distance for all mice;
- movement_hour_ID#.pdf: Individual plots of hourly travel distance for each mouse;
- inter-individual_distance_hour.pdf: Line plot of hourly inter-individual distances for all mouse pairs;
- inter-individual_distance_hour_ID#_vs_ID#.pdf: Individual plots of hourly inter-individual distance for each pair;
- ccr_hour.pdf: Line plot of hourly CCR for all mouse pairs;
- ccr_hour_ID#_vs_ID#.pdf: Individual plots of hourly CCR for each pair.

The R packages used for feature extraction and visualization included tidyverse, lubridate, readxl, and writexl, as listed in **Supplementary Table 3**.

#### S1.4. Zeitgeber time (ZT) conversion

To facilitate circadian-phase-based comparisons, hourly behavioral metrics were converted to Zeitgeber Time (ZT) labels using custom R scripts (IP_HourToZT_Converter_HM.R, IP_HourToZT_Converter_SD.R, and IP_HourToZT_Converter_CCR.R; **Supplementary Table 2**). ZT0 was defined as the onset of the light phase (e.g., 08:00 a.m.). For each hourly time point, the elapsed hours since ZT0 were calculated, and ZT labels were assigned in a 24-hour cycle (ZT0 to ZT23). For example, timepoints were labeled as "Day 1 ZT0", "Day 1 ZT12", etc., to align with the light/dark cycle. Hourly travel distance data were converted using IP_HourToZT_Converter_HM.R (**Supplementary Table2**), and the output (ZT_converted_HM.xlsx) was used for circadian-phase-based analysis of locomotor activity.

Similarly, hourly inter-individual distances were converted using IP_HourToZT_Converter_SD.R (**Supplementary Table2**), with the output (ZT_converted_SD.xlsx) used for phase-based analysis of inter-individual distance. Likewise, hourly CCR values were converted using IP_HourToZT_Converter_CCR.R (**Supplementary Table 2**), with the output (ZT_converted_CCR.xlsx) used for phase-based analysis of social proximity.

#### S1.5. Visualization of ZT-converted behavioral data

ZT-converted behavioral data were visualized using dedicated R scripts (IP_ZT_individual_movement_plot.R, IP_ZT_pairwise_inter-individual_dist_plot.R, IP_ZT_pairwise_CCR_plot.R; **Supplementary Table 2**).

Individual travel distance was visualized as ZT-based time series (Movement_ZT_<ID>.pdf) and as Light/Dark summed plots (Movement_LightDarkSum_<ID>.pdf). Group-level summary plots were also generated (Movement_ZT_All.pdf, Movement_LightDarkSum_All.pdf).

Pairwise inter-individual distance and CCR were similarly visualized for each mouse pair (Inter-individual_distance_ZT_<Pair>.pdf, CCR_ZT_<Pair>.pdf), and Light/Dark summaries were also provided (Inter-individual distance_lightdarkmean_<Pair>.pdf, CCR_LightDarkMean_<Pair>.pdf).

**S1.6. Statistical comparison of two groups**

Group differences (e.g., sex differences) in locomotor activity, inter-individual distance, and CCR were evaluated using R scripts IP_group_comparison_movement.R, IP_group_comparison_inter-individual_dist.R, and IP_group_comparison_CCR.R (**Supplementary Table 2**). For each metric, two-way ANOVA and Tukey's *post hoc* tests were performed for each ZT hour and 12-hour Light/Dark period. Significance was annotated in plots as *$p < 0.05$, **$p < 0.01$, ***$p < 0.001$.

Outputs included:

- Hourly group comparison plots: (HourlyMovement_GroupComparison_.pdf, Inter-individual_dist_group_comparison_.pdf, CCR_GroupComparison_.pdf)
- Light/Dark summed comparison plots: (Sum12h_GroupComparison_.pdf)
- Statistical tables: ANOVA_Hourly_byZT_.csv, Tukey_Hourly_byZT_.csv, ANOVA_Sum12h_byPeriod_.csv, Tukey_Sum12h_byPeriod_.csv, ANOVA_Global_1hr_.csv, ANOVA_Global_12hr_.csv

**S1.7. Radar chart visualization**

Radar charts were generated to provide a compact visualization of multidimensional behavioral profiles and to support intuitive comparisons across sexes and recording days. Radar-chart export was implemented in R using a dedicated script (Radar chart batch exporter; **Supplementary Table 2**), which interactively imports a summary Excel file and outputs PDF files.

The script reads an Excel table annotated by Sex, ID, Day, and light/dark phase (LD), together with three behavioral metrics: travel distance, inter-individual distance, and CCR. The data were reshaped into a wide-format table per mouse and day, yielding six parameters: travel distance, inter-individual distance, and CCR, each quantified for light and dark phases. When multiple entries were present for the same mouse/day/phase, values were aggregated by taking the mean prior to visualization.

Radar charts were exported at both individual and group levels. For individual-level summaries, values were averaged across Days 1–4 to generate one radar chart per mouse, enabling visualization of inter-individual variability within each sex. In addition, day-by-day radar charts were generated for each mouse to visualize temporal changes in behavioral profiles across the recording period. For group-level comparisons, mean profiles were computed by averaging across individuals within each sex, and radar charts were generated for each day (Day 1–4) as well as for the overall average across days.

For sex comparisons in group-level radar charts, statistical differences between males and females were evaluated for each of the six variables using Welch's two-sample *t*-test (two-sided). Significance was annotated on the corresponding axes as *$p < 0.05$, **$p < 0.01$, and ***$p < 0.001$.

**Output files**

The following output files related to radar-chart visualization were generated for each experimental session:

- 00_Individual_ID_DayAvg_raw/:
  Folder containing individual-level radar charts averaged across Days 1–4, separated by sex.
- 00b_Individual_ID_ByDay_raw/:
  Folder containing day-by-day radar charts for each individual mouse (Day 1–Day 4), separated by sex.
- 01_DayAvg_raw_rmax2_Male_vs_Female.pdf:
  Group-level radar chart comparing male and female mean profiles averaged across Days 1–4.
- 01b_DayAvg_raw_SingleSex/:
  Folder containing group-level radar charts for each sex plotted separately.
- 02_DayX_raw_rmax2_Male_vs_Female.pdf (X = 1–4):
  Group-level radar charts comparing male and female profiles for each recording day.

**Software environment**

Data import and reshaping relied on the following R packages: readxl, dplyr, tidyr, and stringr. Radar charts were rendered using base R graphics, and statistical annotations were generated using built-in statistical functions. All packages used for radar-chart generation and visualization are listed in **Supplementary Table 3**.

### Supp. Fig. 1

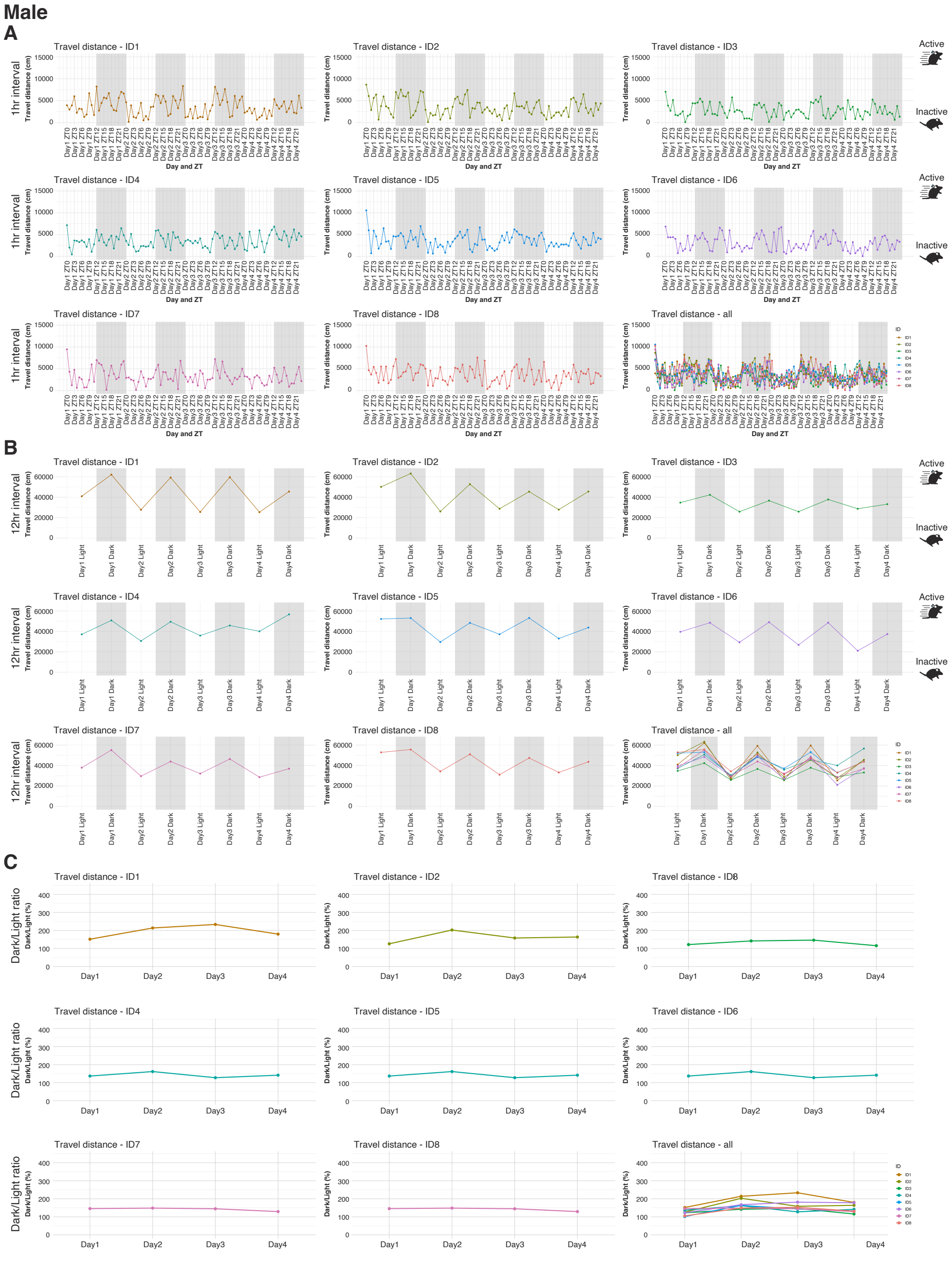


**Supplementary Figure 1. Temporal patterns of travel distance in individual male mice analyzed with the IntelliProfiler 2.0 pipeline.**

(A-C) Travel distance is shown at 1-hour resolution (A), in 12-hour bins (B), and as the dark/light ratio for each day (C). Each panel shows individual trajectories together with the group mean. This figure provides the complete set of individual male mouse plots (n=8) corresponding to in **Fig. 3 A-F**. Abbreviation: ZT, zeitgeber time.

### Supp. Fig. 2

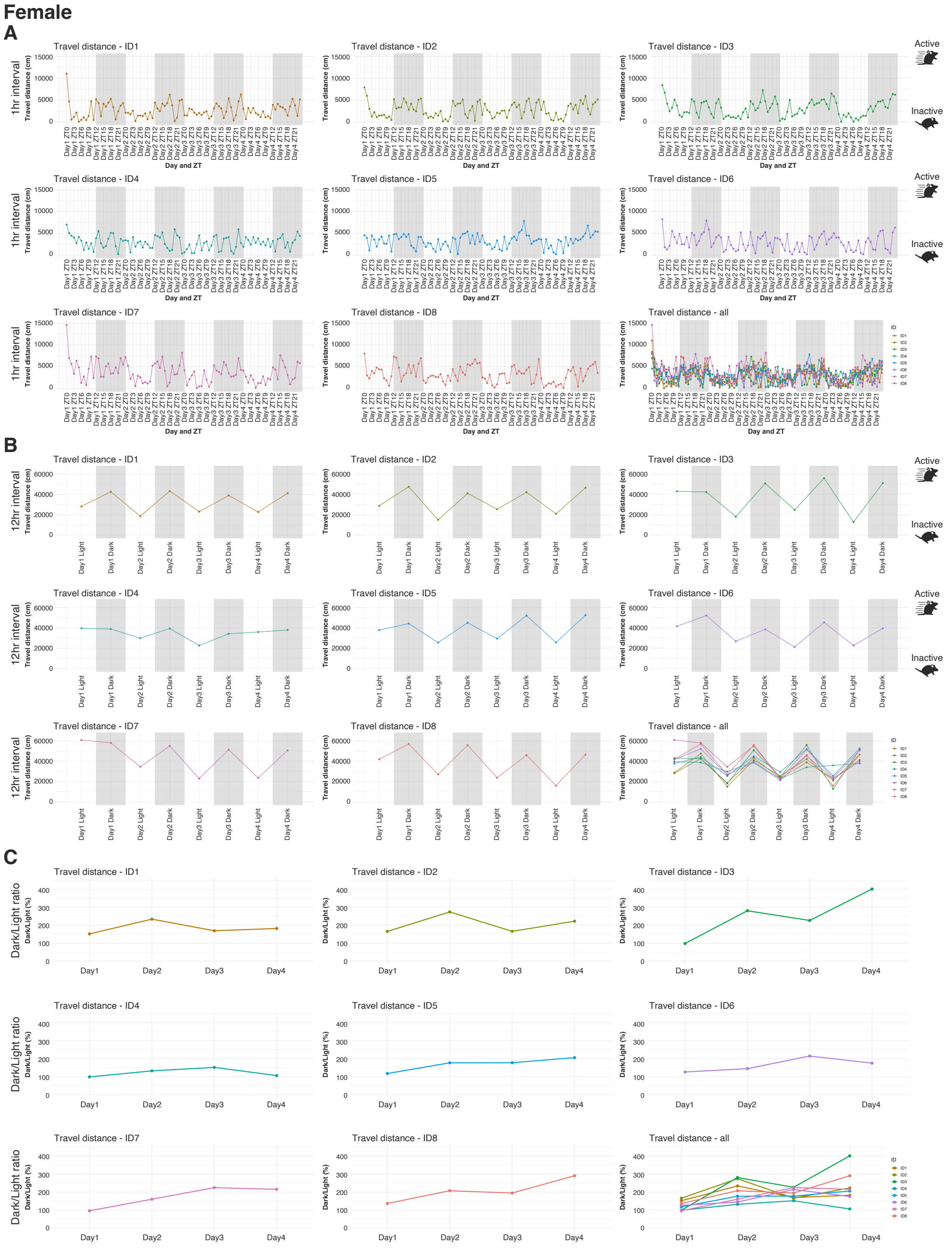


**Supplementary Figure 2. Temporal patterns of travel distance in individual female mice analyzed with the IntelliProfiler 2.0 pipeline.**

(A-C) Travel distance is shown at 1-hour resolution (A), in 12-hour bins (B), and as the dark/light ratio for each day (C). Each panel shows individual trajectories together with the group mean. This figure provides the complete set of individual female mouse plots (n=8) corresponding to in **Fig. 3 A-F**. Abbreviation: ZT, zeitgeber time.

**Supp. Fig. 3**

**Male**

**A**

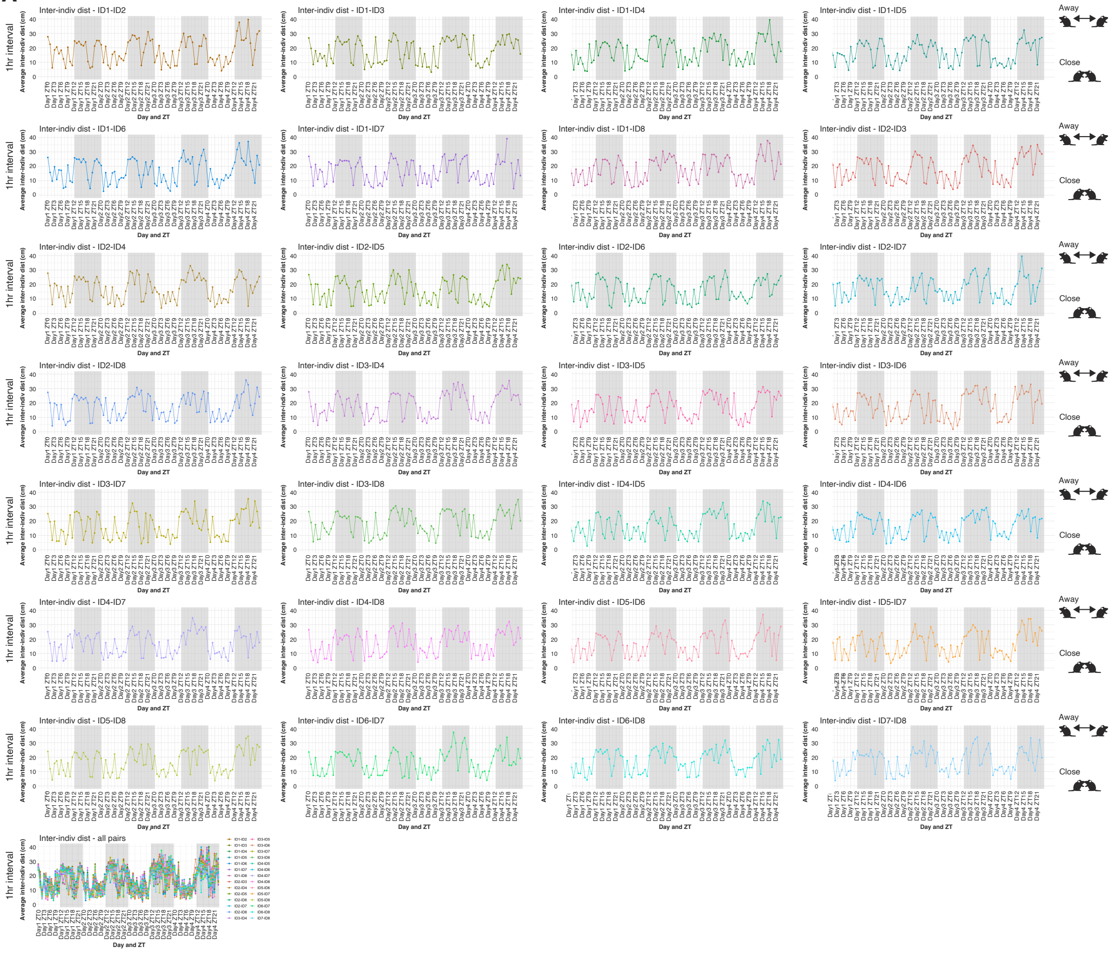


**Supplementary Figure 3. Temporal patterns of inter-individual distance between male mouse pairs analyzed with the IntelliProfiler 2.0 pipeline.**

(A-C) Mean inter-individual distance is shown at 1-hour resolution (A), in 12-hour bins (B), and as the dark/light ratio for each day (C). Each panel shows individual pairwise trajectories together with overall mean across all pairs (n=8; 28 possible pair combination). Two representative male mouse pairs are shown in the corresponding panels in **Fig. 4 A-F**. Abbreviation: ZT, zeitgeber time.

### Supp. Fig. 3
#### Male
### B

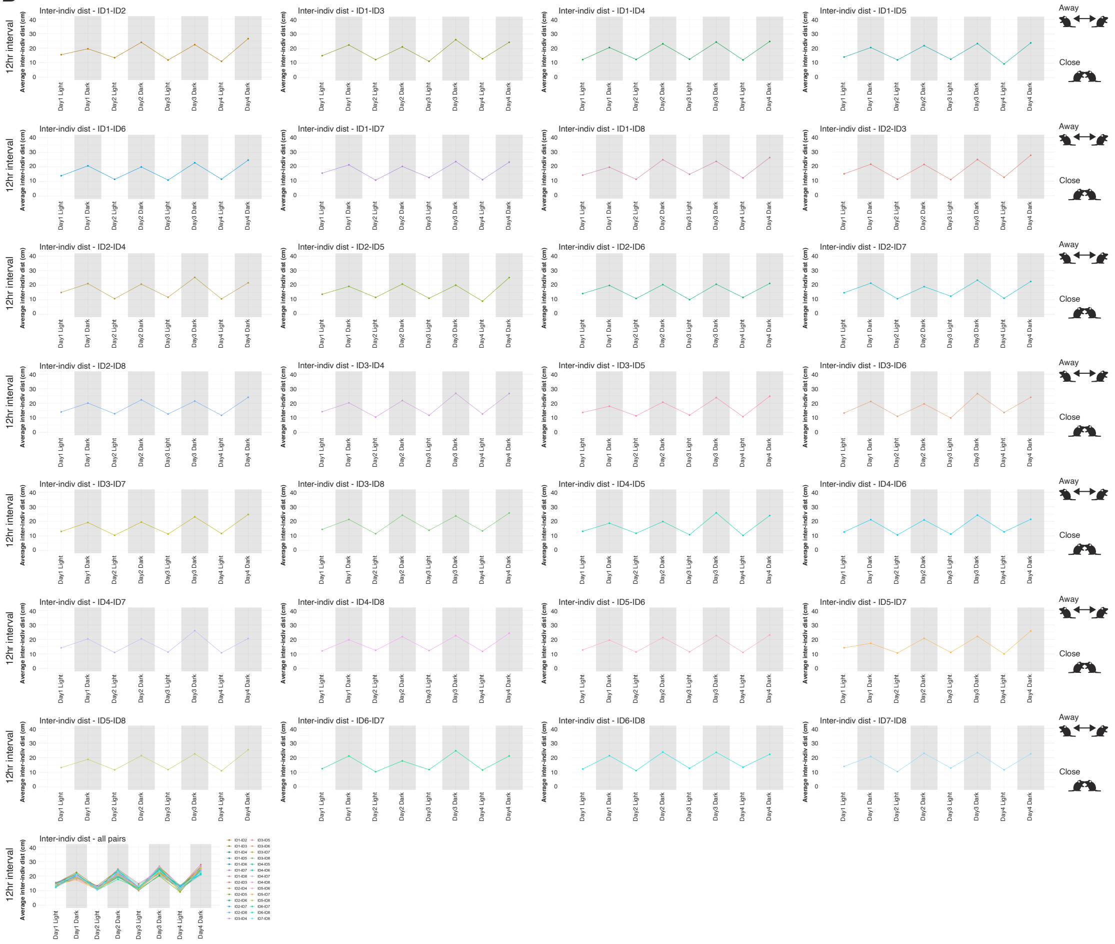


**Supplementary Figure 3** continued.

**Supp. Fig. 3**

**Male**

**C**

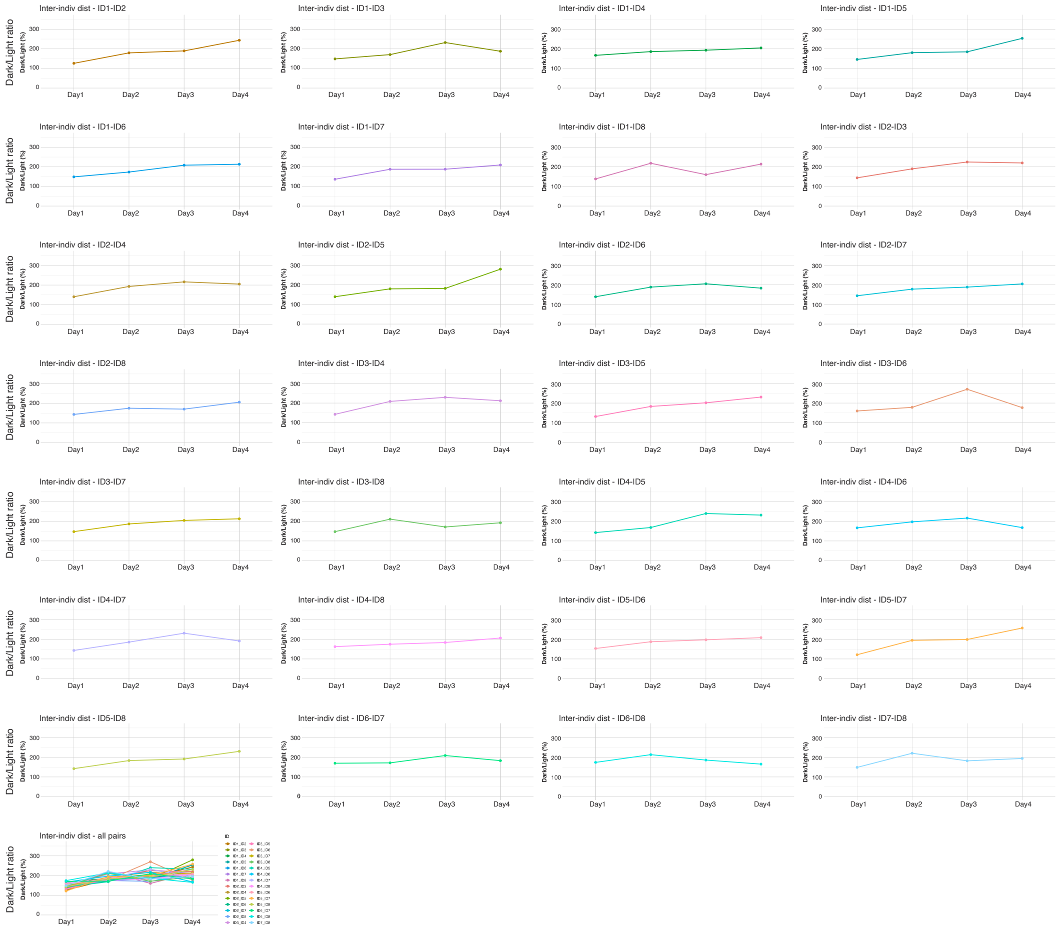


**Supplementary Figure 3** continued.

### Supp. Fig. 4

#### Female

### A

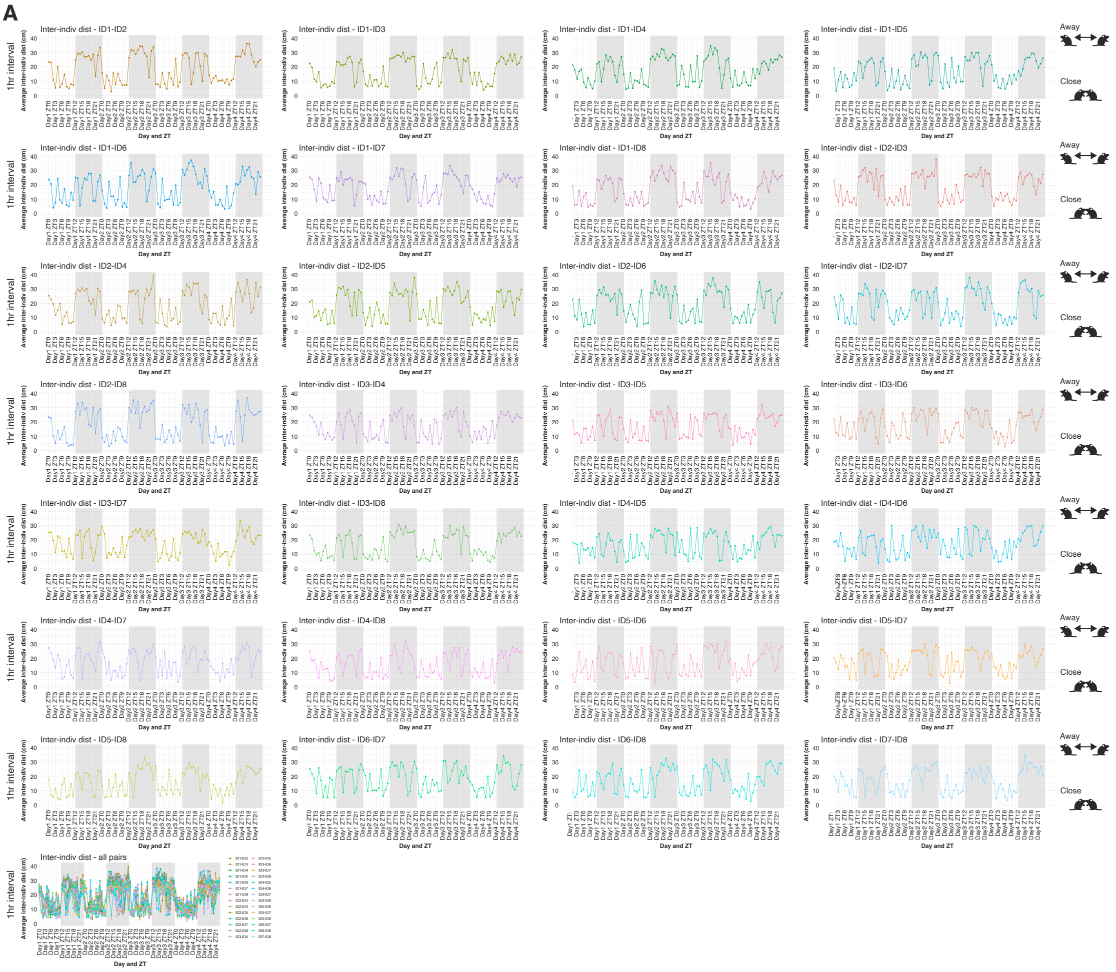


**Supplementary Figure 4. Temporal patterns of inter-individual distance between female mouse pairs using the IntelliProfiler 2.0 pipeline.**

(A-C) Mean inter-individual distance is shown at 1-hour resolution (A), in 12-hour bins (B), and as the dark/light ratio for each day (C). Each panel shows individual pairwise trajectories together with overall mean across all pairs (n=8; 28 possible pair combination). Two representative male mouse pairs are shown in the corresponding panels in **Fig. 4 A-F**. Abbreviation: ZT, zeitgeber time.

**Supp. Fig. 4**

**Male**

**B**

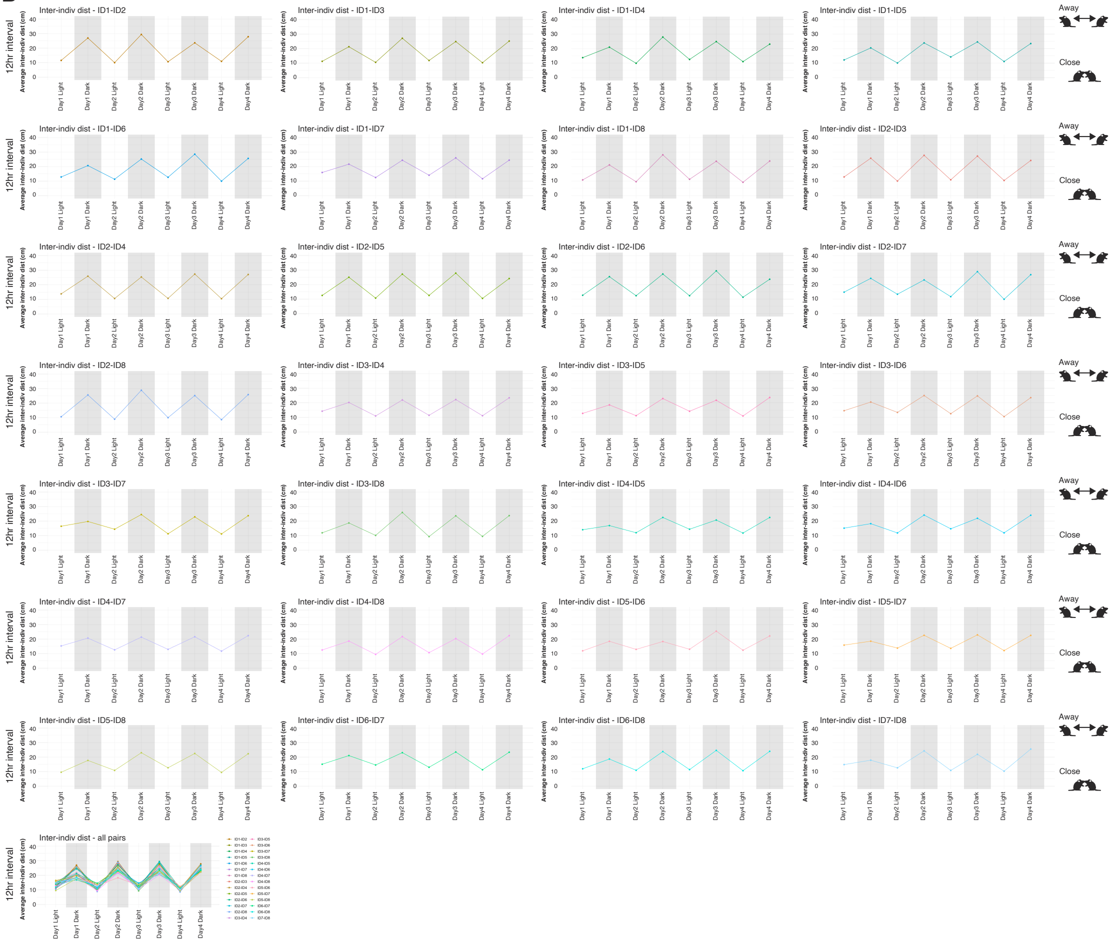

**Supplementary Figure 4** continued.

### Supp. Fig. 4

#### Female

### C

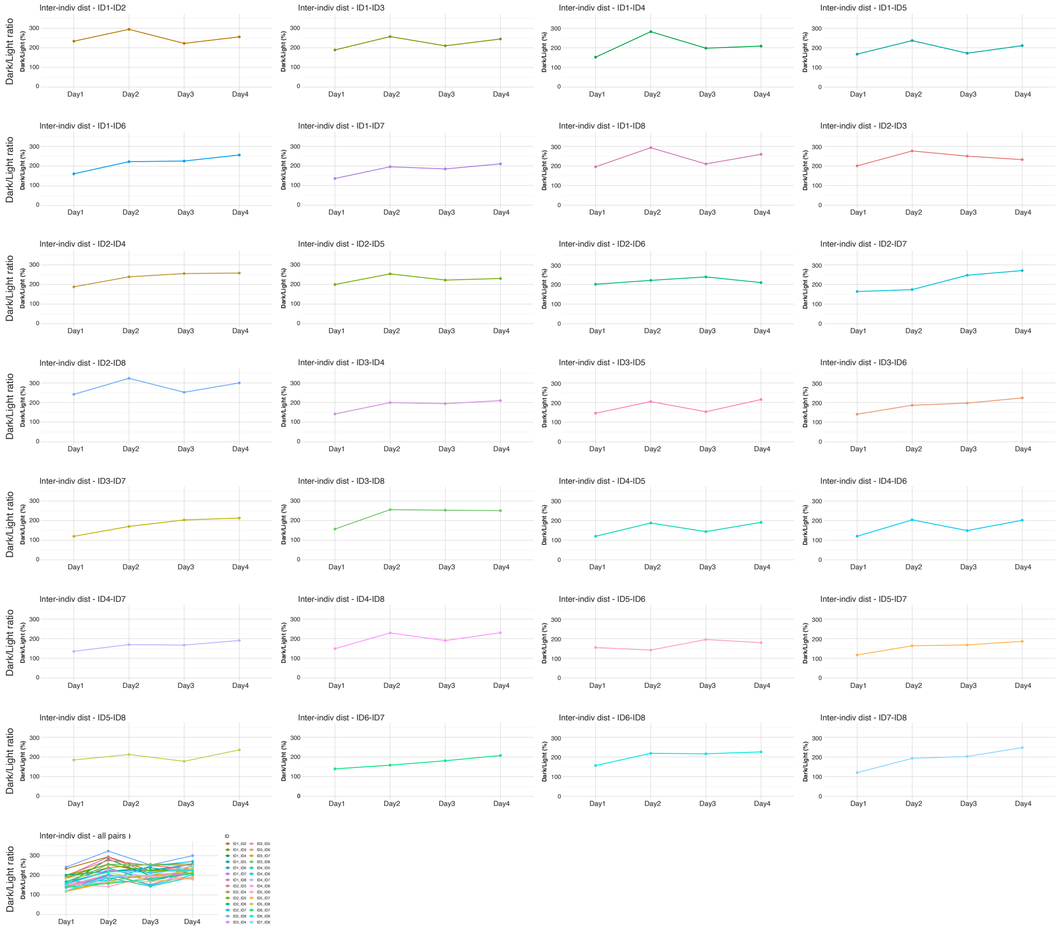


**Supplementary Figure 4** continued.

### Supp. Fig. 5

#### Male

### A

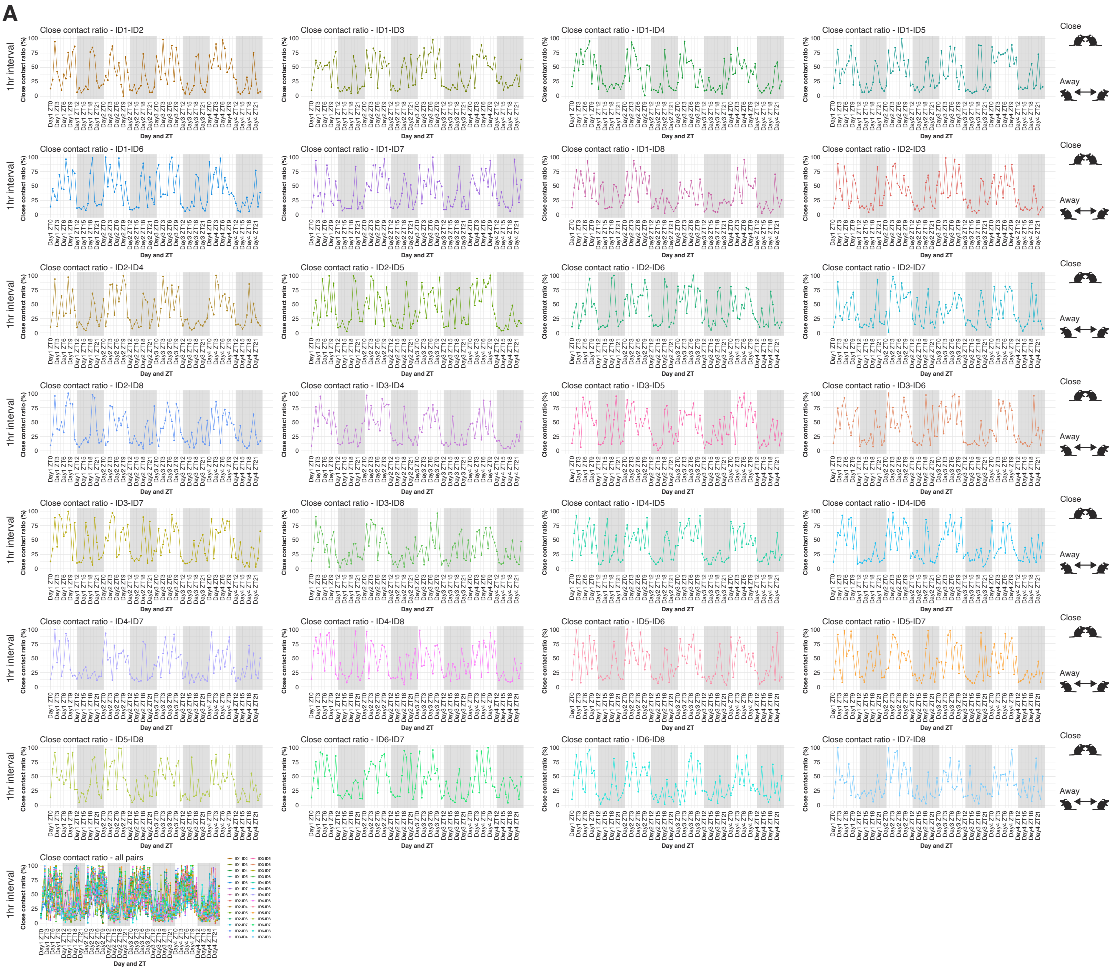


**Supplementary Figure 5. Temporal patterns of close contact ratio (CCR) between male mouse pairs analyzed with the IntelliProfiler 2.0 pipeline.**

(A-C) CCR (percentage of time in close contact) is shown at 1-hour resolution (A), 12-hour bins (B), and as the dark/light ratio for each day (C). Each panel shows individual pairwise trajectories together with the overall mean across all pairs (n=8; 28 possible pair combinations). Two representative male mouse pairs are shown in the corresponding panels in **Fig. 5 B-G**. Abbreviation: ZT, zeitgeber time.

### Supp. Fig. 5
#### Male
### B

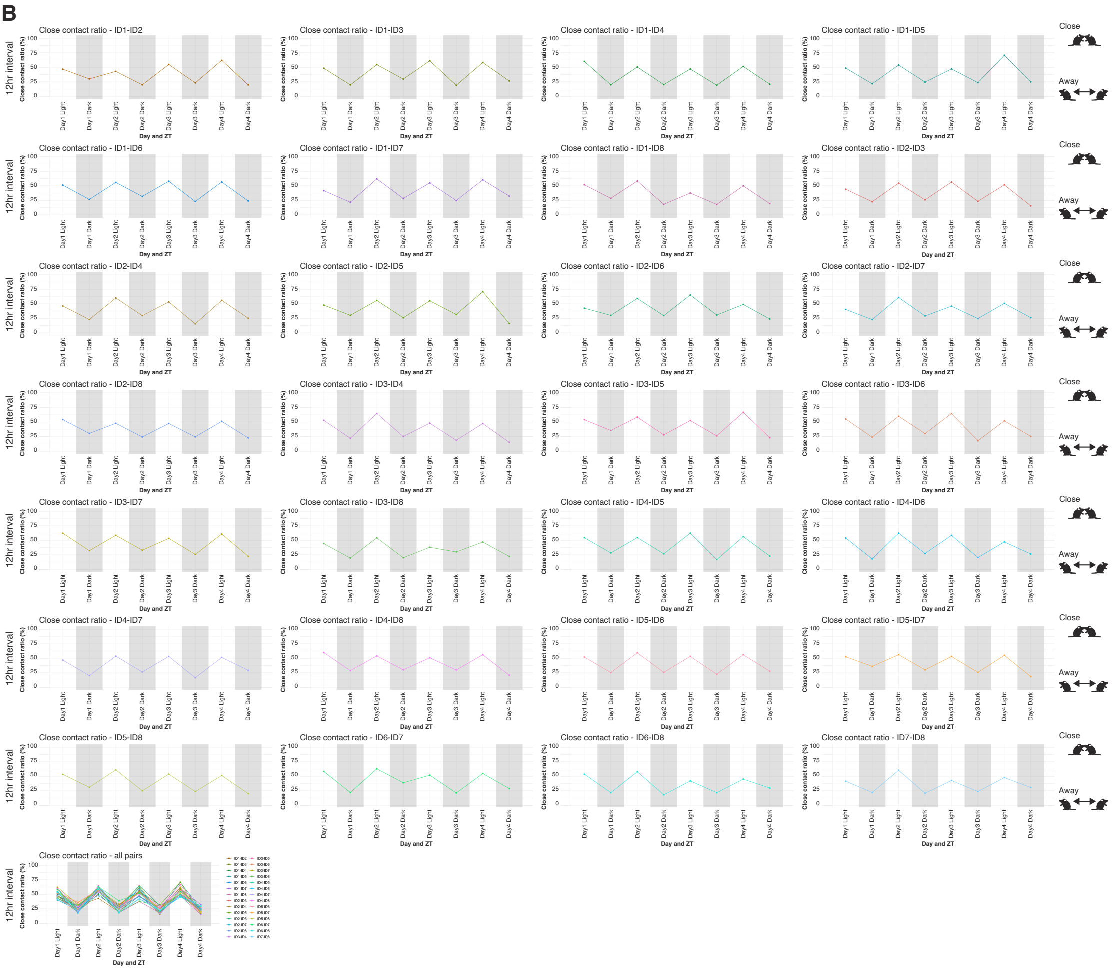


**Supplementary Figure 5** continued.

### Supp. Fig. 5
### Male
## C

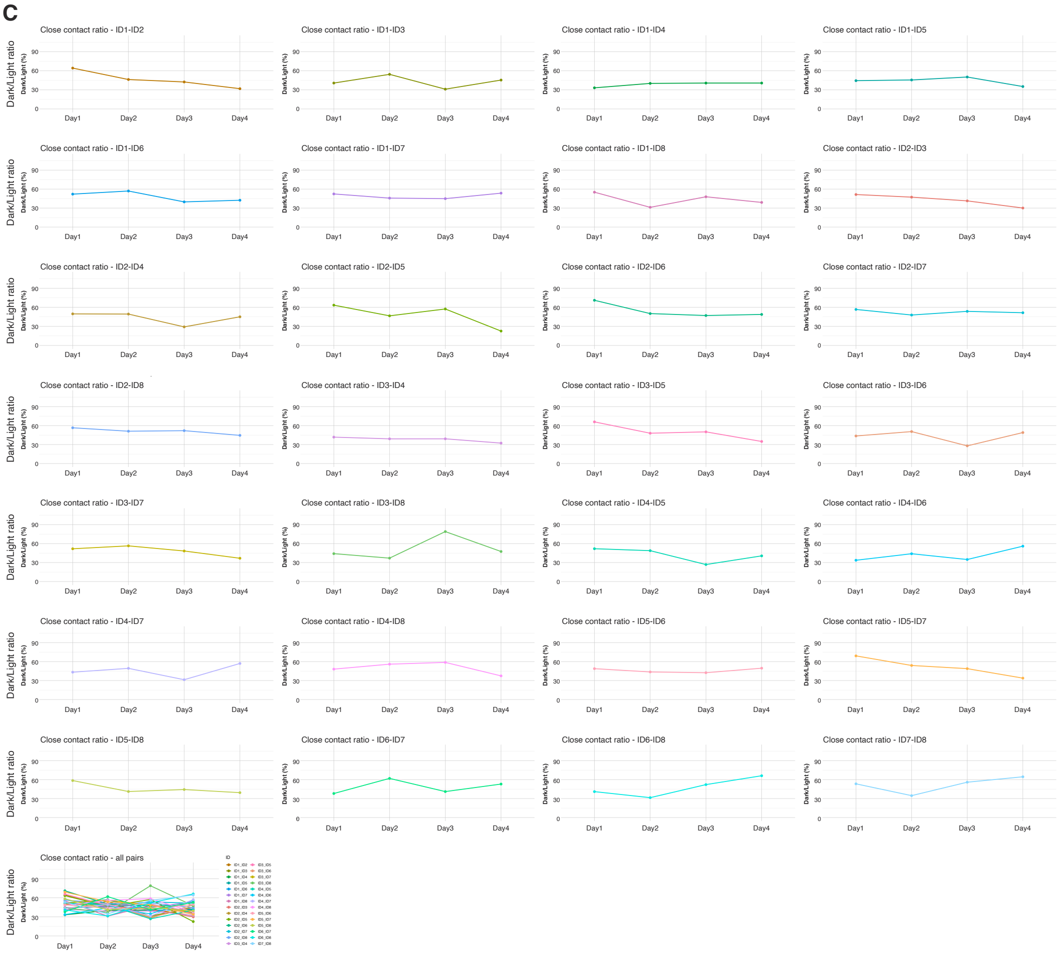


**Supplementary Figure 5** continued.

### Supp. Fig. 6

#### Female

### A

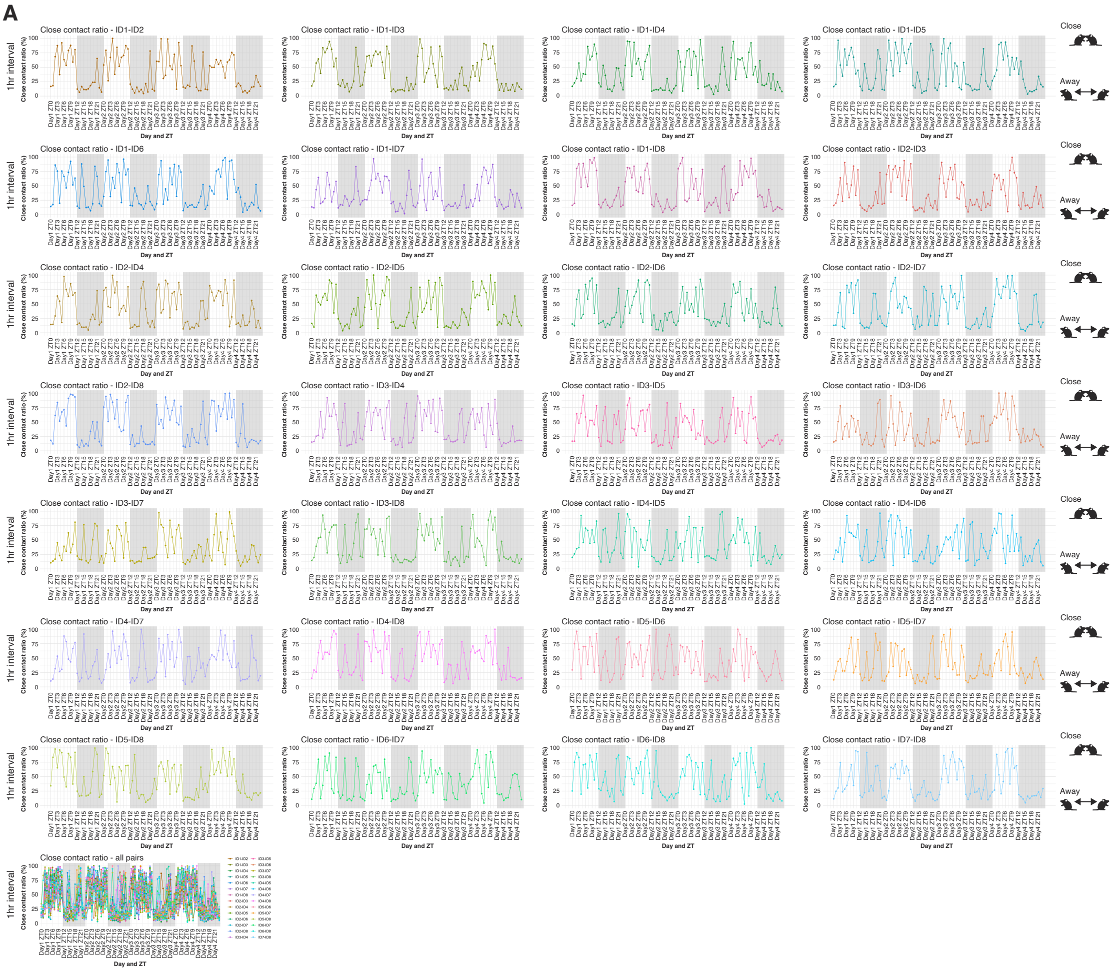


**Supplementary Figure 6. Temporal patterns of close contact ratio (CCR) between female mouse pairs analyzed with the IntelliProfiler 2.0 pipeline.**

(A-C) CCR (percentage of time in close contact) is shown at 1-hour resolution (A), 12-hour bins (B), and as the dark/light ratio for each day (C). Each panel shows individual pairwise trajectories together with the overall mean across all pairs (n=8; 28 possible pair combinations). Two representative female mouse pairs are shown in the corresponding panels in **Fig. 5 B-G**. Abbreviation: ZT, zeitgeber time.

### Supp. Fig. 6

#### Female

### B

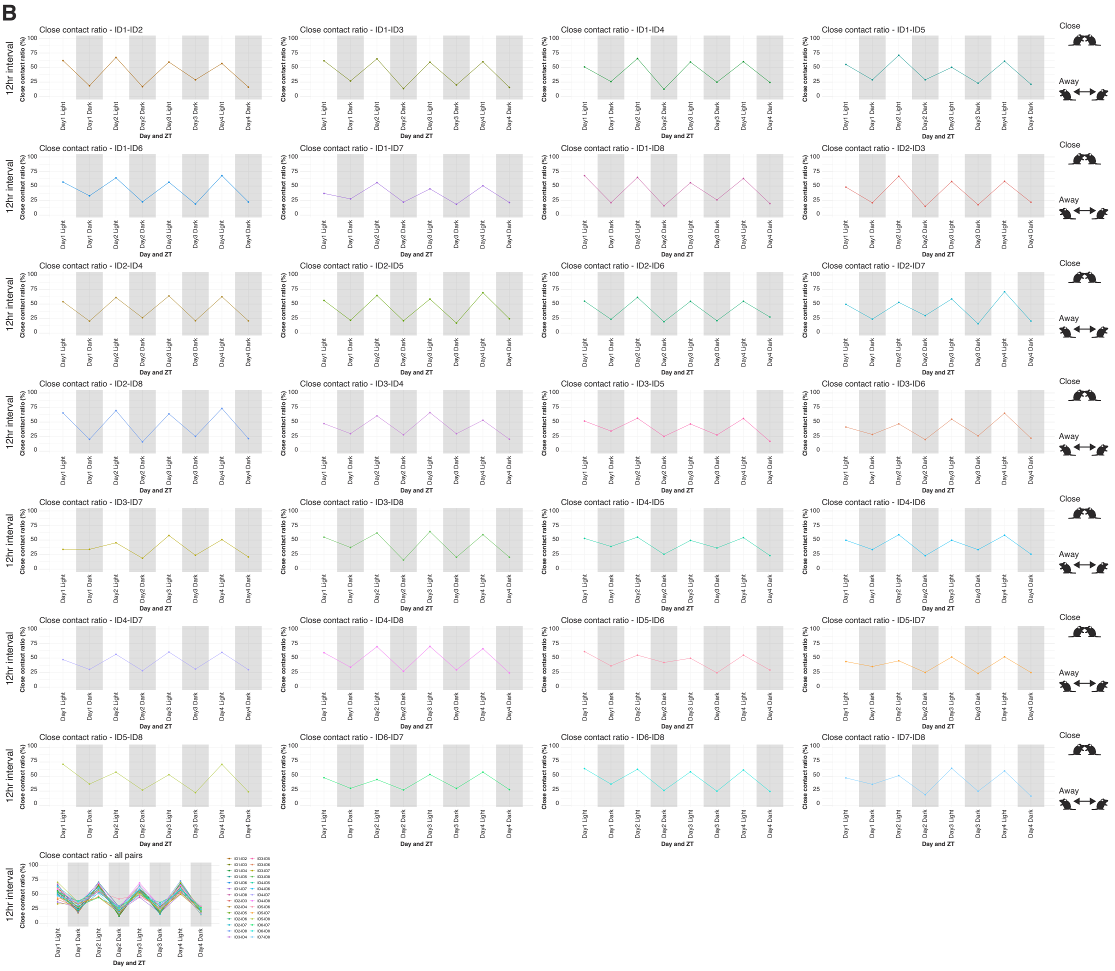


**Supplementary Figure 6** continued.

### Supp. Fig. 6
#### Female
### C

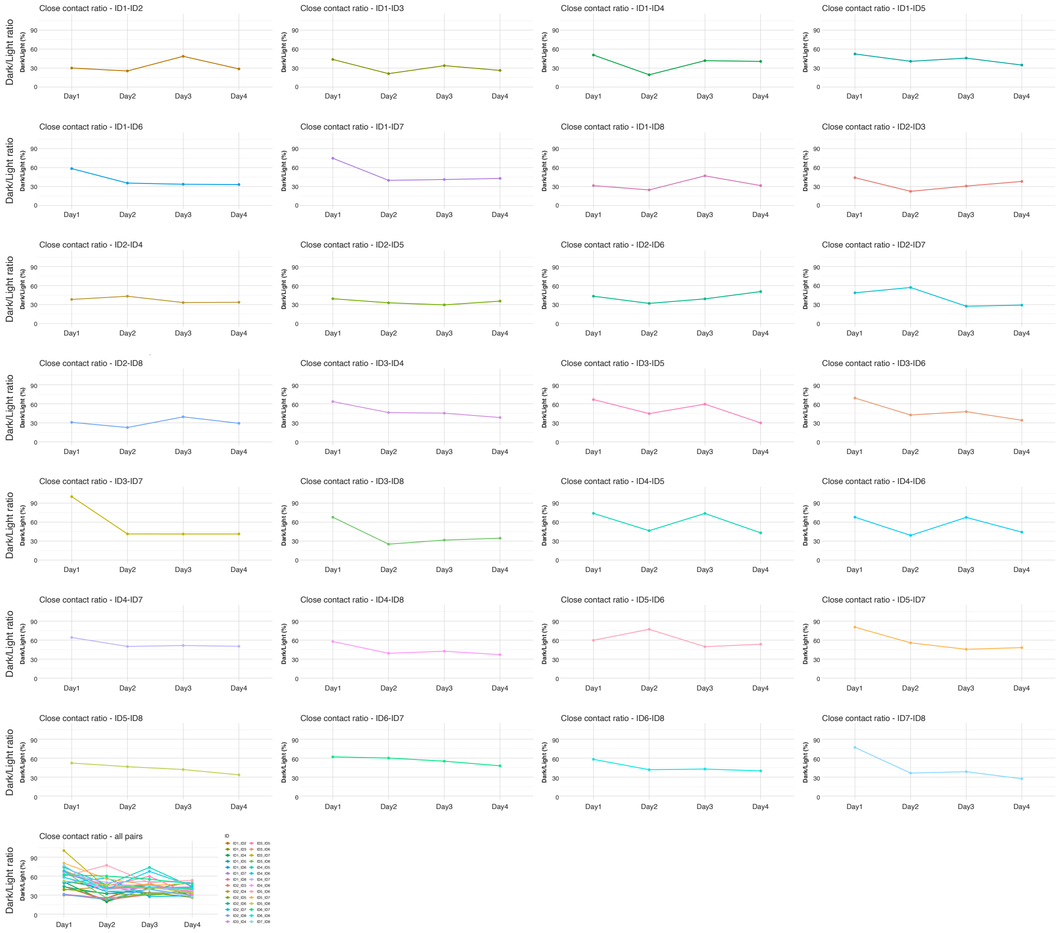


**Supplementary Figure 6** continued.

**Supp. Fig. 7**
**Male**
**A**

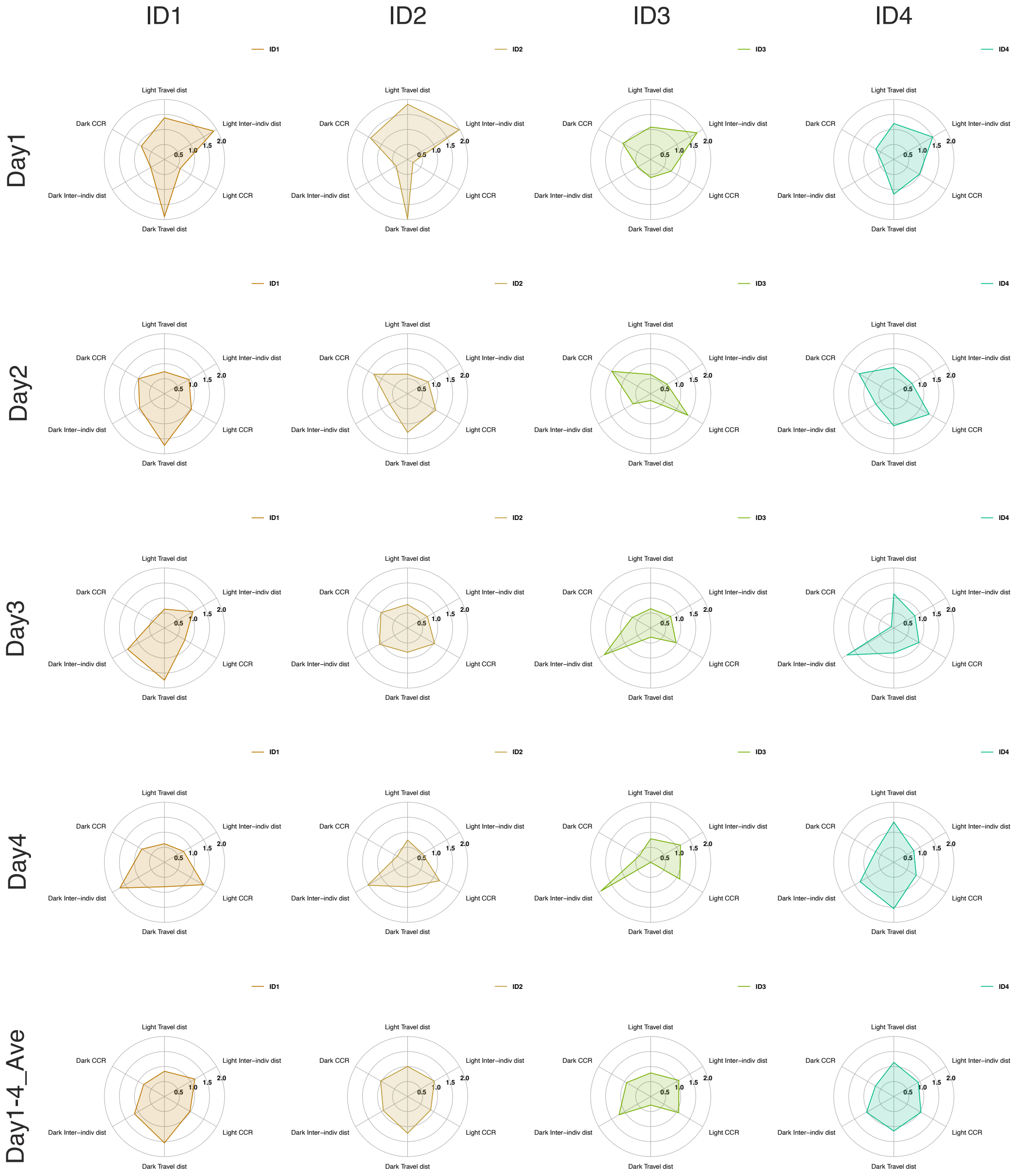


**Supplementary Figure 7. Radar-chart visualization of multidimensional behavioral profiles across recording days.**
(A, B) Radar charts summarize the mean behavioral profile of male (A) and female (B) groups (n=8 per sex) for each recording day and for the average across Days 1-4. Parameters are defined as travel distance, inter-individual distance, and CCR, each quantified for light and dark phases. Abbreviations; CCR, close contact ratio. Abbreviations; CCR, close contact ratio.

**Supp. Fig. 7**
**Male**
**A**

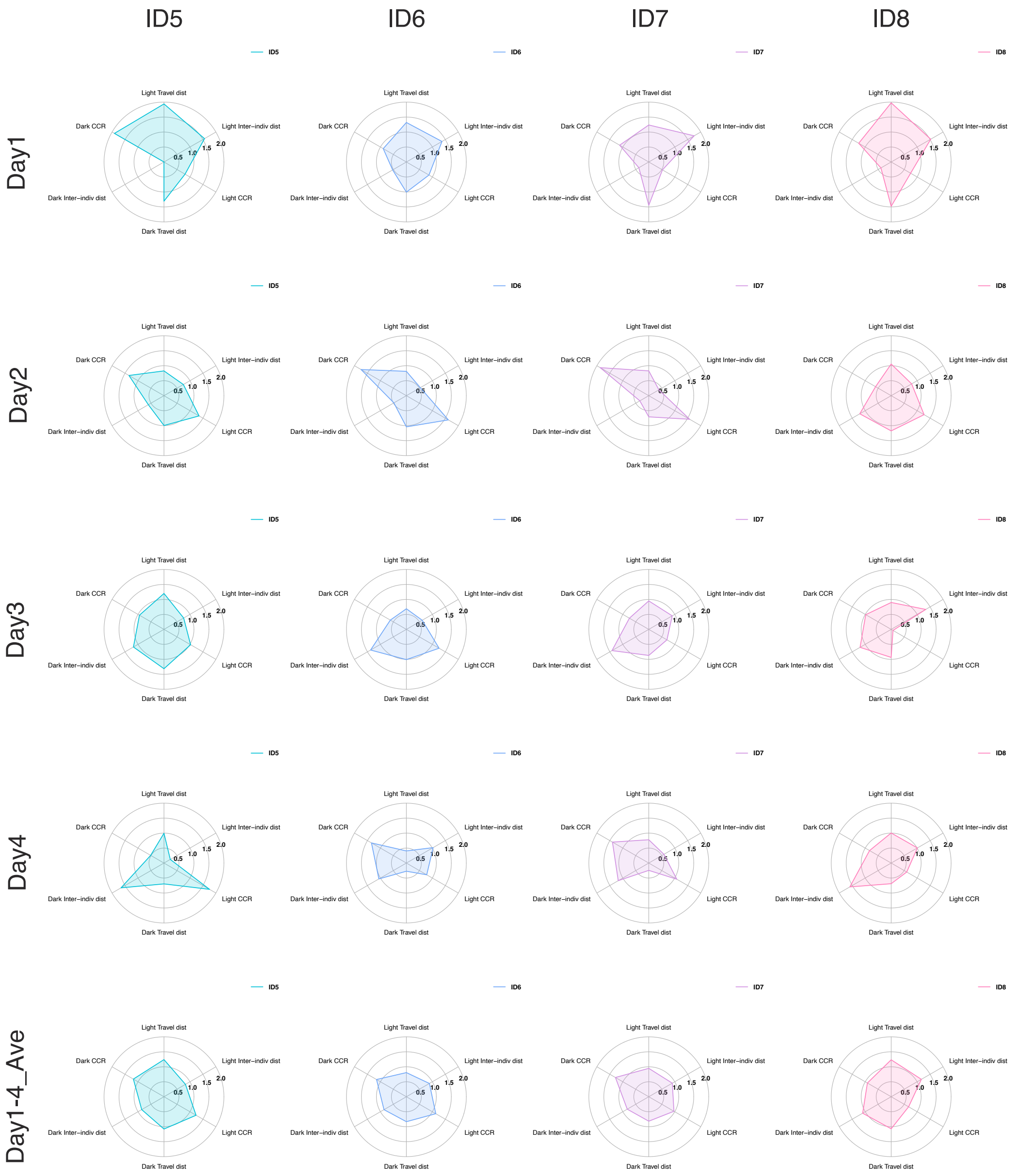


**Supplementary Figure 7** continued.

**Supp. Fig. 7**
**Female**
**B**

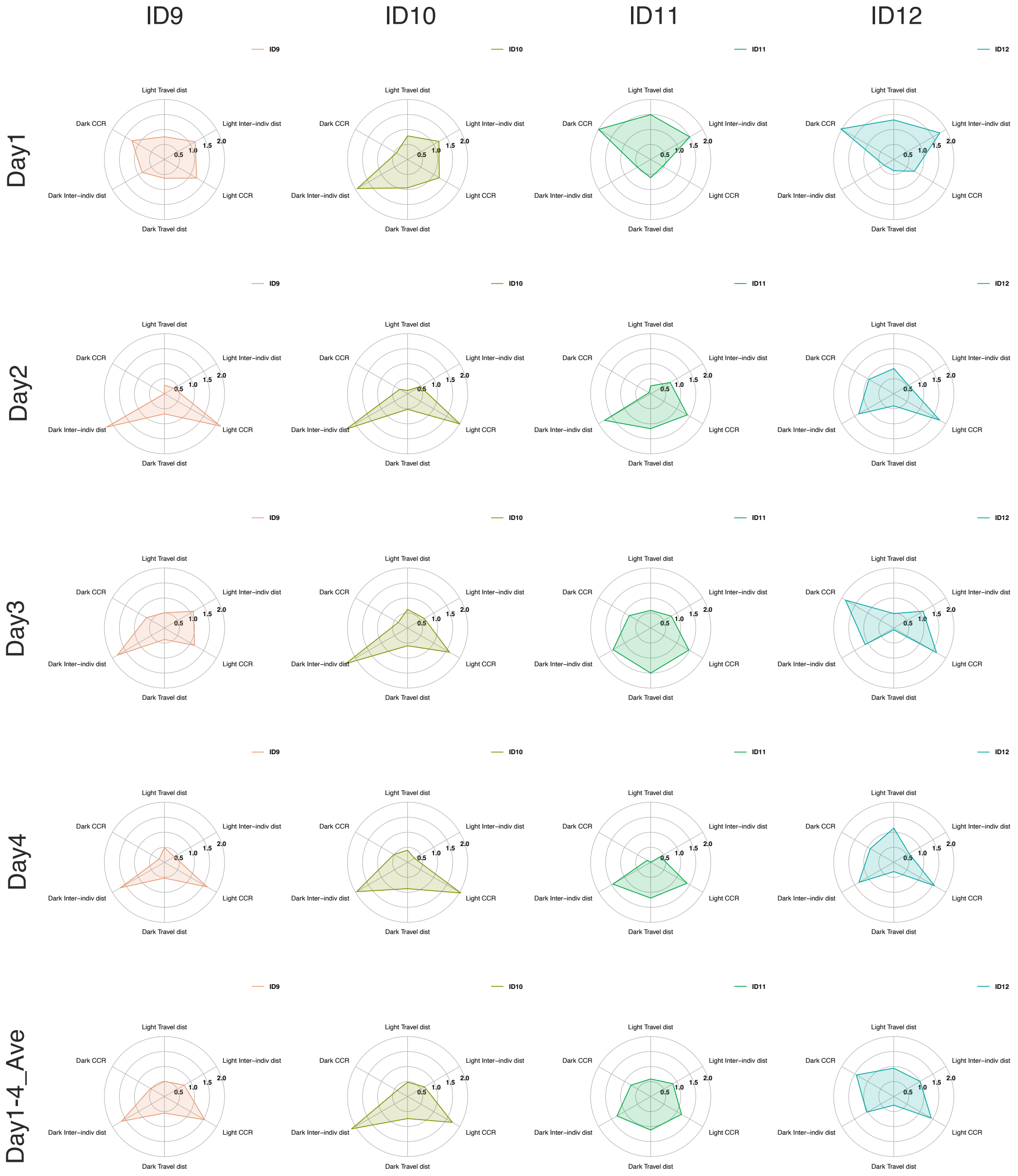


**Supplementary Figure 7** continued.

**Supp. Fig. 7**
**Female**
**B**

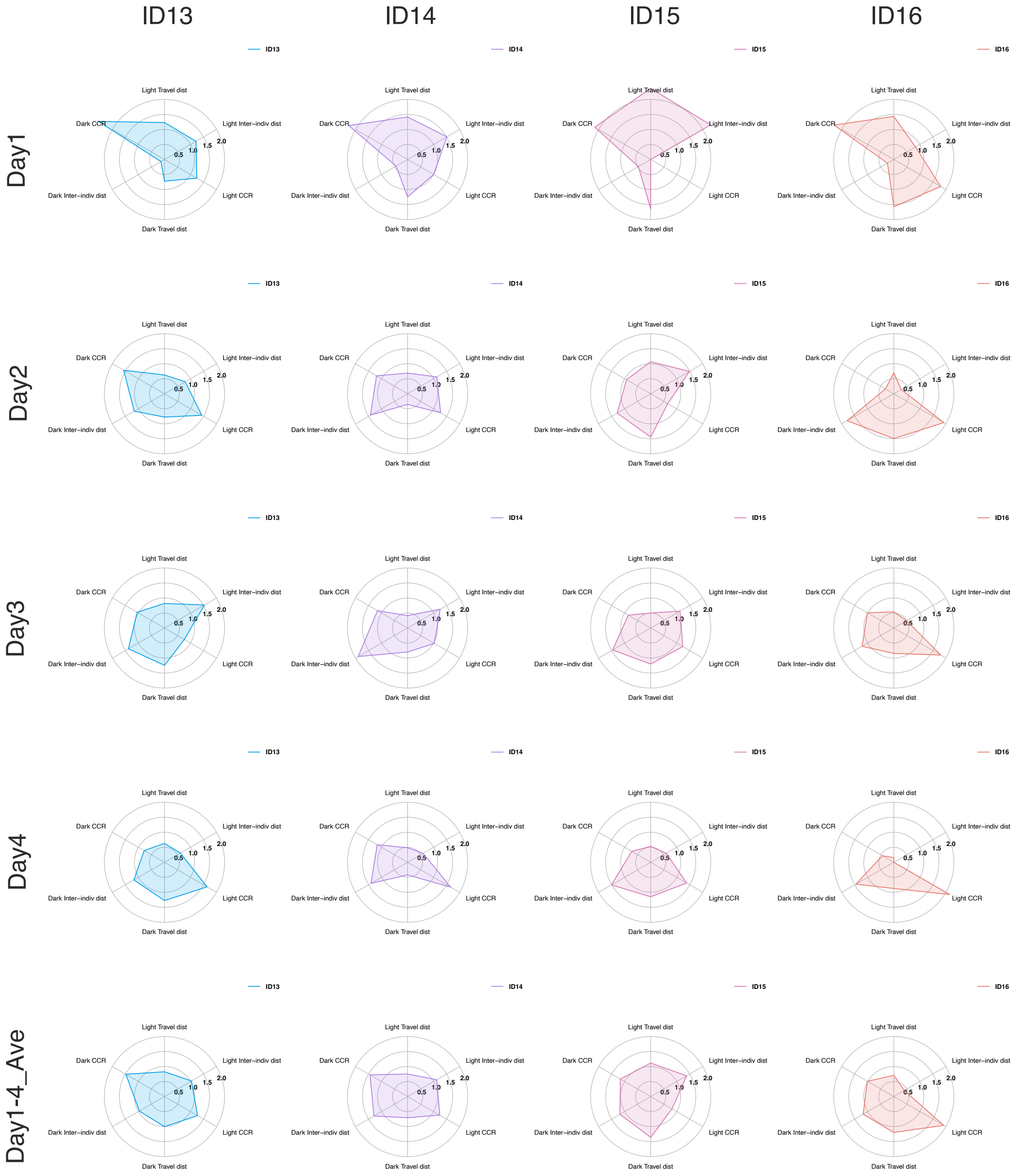


**Supplementary Figure 7** continued.

**Supp. Fig. 8**

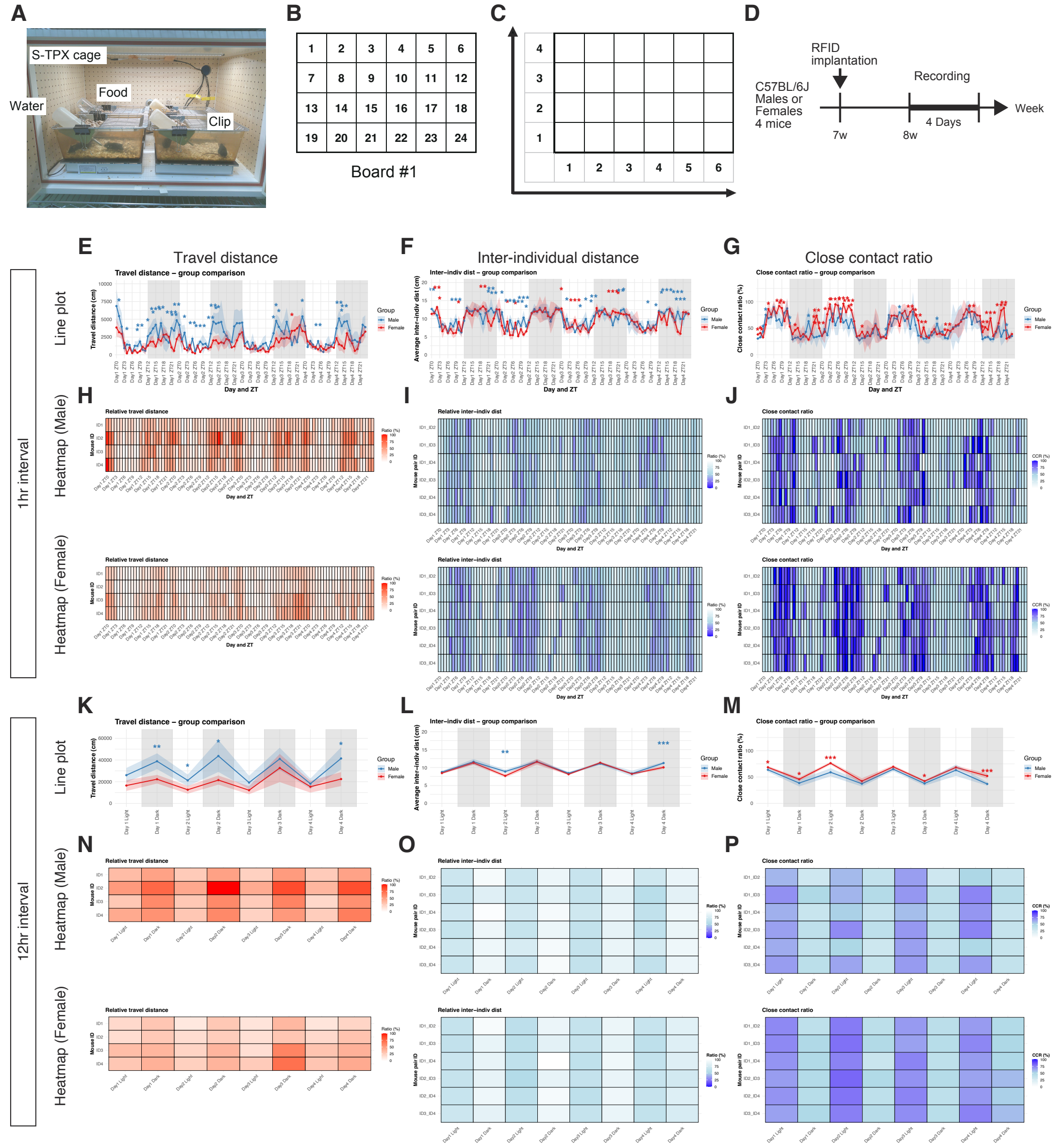


**Supplementary Figure 8. Temporal patterns obtained using a single eeeHive 2D board.** (A) Group behavior recording in a small cage on an eeeHive 2D board. (B) Layout and antenna numbering of an eeeHive 2D board. (C) Conversion of antenna positions to XY coordinates (X1-6, and Y1-8) in IP_single pipeline. (D) Experimental timeline. RFID tags were implanted in 7-week-old C57BL/6 mice and group behavior was recorded for four consecutive days at 8-week-old. (E-P) Comparisons between male and female groups (n=4 per sex) across four consecutive days. Panels show cumulative travel distance (E, H, K, N), mean inter-individual distance (F, I, L, O), and CCR (G, J, M, P) at 1-hour resolution (E-J) and in 12-hour bins (K-P). Line plots are shown in (E-G, K-M), and heatmaps in (H-J, N-P). Statistical significance: $^{*}p < 0.05$, $^{**}p < 0.01$, $^{***}p < 0.001$. Male and female data are shown in blue and red, respectively; shaded ribbons indicate standard deviation (SD). Abbreviations; CCR, close contact ratio.

**Supplementary Table1. Comprehensive list of equipment, materials, and animals used for RFID-based behavioral monitoring.**

The table summarizes the resources used for subcutaneous RFID tag implantation, anesthesia, housing, and behavioral data acquisition using the eeeHive 2D RFID sensor array and IntelliProfiler analysis pipeline.

**Supplementary Table2. List of R scripts used in IntelliProfiler analysis.**

The table describes each script's name, purpose, input files, output files, and R packages used.

**Supplementary Table 3. List of R packages used in IntelliProfiler analysis.**

The table describes the R packages and their versions.

**Supplementary Table4. Example mapping between short IDs and original RFID tag identifiers used for IntelliProfiler analysis.**

The table lists the correspondence between assigned short IDs (ID1–ID8) and the original RFID tag IDs for male mice, as read by the eeeHive 2D RFID sensor array.

**Supplementary Movie 1. Video of RFID tag implantation procedure in a mouse under anesthesia.**
